# How many markers are needed to robustly determine a cell’s type?

**DOI:** 10.1101/2021.04.16.439807

**Authors:** Stephan Fischer, Jesse Gillis

**Affiliations:** Cold Spring Harbor Laboratory, Stanley Institute for Cognitive Genomics, Cold Spring Harbor, NY 11724, USA; Cold Spring Harbor Laboratory, Watson School of Biological Sciences, Cold Spring Harbor, NY, USA

**Keywords:** Cell types, single-cell RNA sequencing, replicability, marker genes, cell type taxonomy

## Abstract

Our understanding of cell types has advanced considerably with the publication of single cell atlases. Marker genes play an essential role for experimental validation and computational analyses such as physiological characterization through pathway enrichment, annotation, and deconvolution. However, a framework for quantifying marker replicability and picking replicable markers is currently lacking. Here, using high quality data from the Brain Initiative Cell Census Network (BICCN), we systematically investigate marker replicability for 85 neuronal cell types. We show that, due to dataset-specific noise, we need to combine 5 datasets to obtain robust differentially expressed (DE) genes, particularly for rare populations and lowly expressed genes. We estimate that 10 to 200 meta-analytic markers provide optimal performance in downstream computational tasks. Replicable marker lists condense single cell atlases into interpretable and generalizable information about cell types, opening avenues for downstream applications, including cell type annotation, selection of gene panels and bulk data deconvolution.

## Introduction

Recent atlas efforts based on single cell technologies have led to comprehensive cell type taxonomies that include a multitude of novel cell types (Tasic et al. 2018; Zeisel et al. 2018; Schaum et al. 2018; Packer et al. 2019; Cao et al. 2020). The discovery of new cell types and novel biological heterogeneity served as a foundation for promising avenues for the understanding of tissue homeostasis and disease. However, to develop downstream applications and experiments, an actionable description of cell types is required that extends beyond taxonomic classification. While sporadic post-hoc markers are published alongside taxonomies, the replicability of these markers is rarely assessed. Here, we systematically evaluate marker replicability and propose unprecedented lists of replicable markers (or meta-markers) for neuronal cell types by selecting an optimal number of robustly upregulated genes across a compendium of brain datasets.

Given the rapid progression in the number and size of single-cell datasets (Svensson et al. 2018), making atlases easily accessible is an increasingly difficult challenge. Cell type centroids provide an efficient summary of active gene expression programs (Zeisel et al. 2018), but they are subject to batch effects (Tung et al. 2017) and discard expression variability. While integrative methods have been successful at mitigating batch effects for the joint analysis of a small groups of datasets (Butler et al. 2018; Haghverdi et al. 2018; Welch et al. 2019; Korsunsky et al. 2019; Lin et al. 2019) and the transfer of cell type annotations (Kiselev et al. 2018; Stuart et al. 2019), the abstract embedding of cell types is costly, as well as difficult to interpret and to extract for downstream applications. In contrast, markers provide an interpretable common denominator that does not involve data re-analysis or complex mathematical transformations; they are commonly used for functional characterization (Mancarci et al. 2017), cell type annotation (Poulin et al. 2016; Johnson and Walsh 2017; Pliner et al. 2019; Zhang et al. 2019), deconvolution of bulk data (Wang et al. 2019; Newman et al. 2019; Patrick et al. 2020) and spatial data (Qian et al. 2020), selection of representative gene panels (Moffitt et al. 2018), cross-species comparisons (Tosches et al. 2018; Hodge et al. 2019; Krienen et al. 2019; Bakken et al. 2020), and mapping of organoids to in vivo progenitors (Velasco et al. 2019; Bhaduri et al. 2020). For many of these applications, the strength of individual markers is limited by the lack of conservation (Bakken et al. 2020) and the sporadic expression in individual cells (Kharchenko et al. 2014; Risso et al. 2018; Hicks et al. 2018; Chen and Zhou 2018). Moving past individual markers to small lists is done sporadically to capture combinatorial relationships or improve power, but has not yet exploited the full power of scRNA-seq data. In specific, because cell types are encoded in a low-dimensional expression space (Crow and Gillis 2018), we hypothesize that they can be captured with high resolution and generalizable definitions using redundant and robust marker lists. These lists can then easily be compared and combined across datasets for downstream analyses.

The problem of finding generalizable descriptions of cell types has a long history in the brain, famously illustrated by Ramon y Cajal’s morphology-based descriptions (RAMON Y CAJAL 1904). More recently, the Petilla convention emphasized the need to describe neurons according to a multi-modal taxonomy, including morphology, electrophysiology, connectivity and transcriptomics (Ascoli et al. 2008). Single cell data, while only covering one aspect of this multi-modal description, have enabled unprecedented wide and deep sampling of brain cells, with current taxonomies containing several hundred cell types (Tasic et al. 2018; Zeisel et al. 2018). They thus offer a chance to assess the robustness of transcriptomic cell types, but current cell types are usually defined based on data from a single lab and a single computational method, while an ideal description should be community-based and method-independent (Yuste et al. 2020). With the recent publication of several single-cell compendia by the Brain Initiative Cell Census Network (BICCN)(Yao et al. 2020a, 2020b), the brain offers a unique opportunity to characterize marker-based descriptions.

In this manuscript, we systematically assess the replicability of markers for BICCN cell types. We identify robust markers (meta-markers) across a compendium of 7 brain single cell datasets containing a total of 482,712 cells from the BICCN, one of the most complex and comprehensive cell type taxonomies to date. The assessment procedure is based on two simple steps: (i) identify markers from single datasets, (ii) obtain a list of meta-markers by selecting replicable markers. The compendium samples from 6 single-cell and single-nuclei technologies, resulting in meta-markers that are robust to the varying sensitivity and contamination levels of these technologies. We further investigate the ability of markers to recapitulate cell types at various levels of granularity. We define two simple performance axes, intuitively representing coverage (being expressed in all cells of interest) and signal-to-noise ratio (being expressed exclusively in cells of interest), that can be efficiently summarized using standard differential expression statistics. While individual meta-markers only imperfectly capture cell types, we find that aggregating 10 to 200 meta-markers leads to optimal performance in downstream computational analyses, such as cell type annotation and deconvolution. Remarkably, these marker-based descriptions, derived from the primary motor cortex, generalize to other cortical brain regions, enabling accurate annotation of individual cells. Robust meta-markers thus provide a simple and actionable description of BICCN cell types, which we make available as high-quality marker lists (Sup. Data 1-3) ranging from the lowest resolution (excitatory neurons, inhibitory neurons, non-neurons) to the finest resolution defined by the BICCN (85 neuronal cell types).

## Results

The ideal marker gene fulfills two criteria: (1) it is expressed in all cells of the population of interest, providing high coverage, (2) it is not expressed in background cells, providing a high signal-to-noise ratio (Fig. 1a). In recently published atlases, it is often unclear how strongly and robustly the proposed markers fulfill these criteria, particularly at high clustering resolution. To investigate replicability of marker strength, our basic strategy is to look for simple statistics that can be robustly averaged across datasets and correctly capture coverage and signal-to-noise. We focused on a BICCN neuron atlas containing 7 datasets with 482,712 cells, organized into a hierarchy of 116 cell types in 3 levels of increasing resolution: classes, subclasses and clusters (Yao et al. 2020a)(Table 1).

**Table 1.** List of Brain Initiative Cell Census Network (BICCN) datasets used in this study. All datasets are from mouse. MOp corresponds to the primary motor cortex, AUD to the auditory cortex. The “# genes detected” column contains the median number of genes detected per cell. The “# UMI / reads” column contains either the median number of reads per cell (for SmartSeq datasets) or the median number of UMIs per cell (for 10X datasets).

| Dataset | Brain regions | Assay | Technology | # cells | # cell types | # genes detected | # UMIs / reads |
| --- | --- | --- | --- | --- | --- | --- | --- |
| scSS | MOp | Cell | SmartSeq | 6,288 | 61 | 9,420 | 1,750,664 |
| snSS | MOp | Nucleus | SmartSeq | 6,171 | 46 | 4,363 | 613,762 |
| scCv2 | MOp | Cell | 10X v2 | 122,641 | 90 | 4,594 | 12,697 |
| snCv2 | MOp | Nucleus | 10X v2 | 76,525 | 43 | 1,716 | 3,145 |
| snCv3M | MOp | Nucleus | 10X v3 | 159,738 | 113 | 4,237 | 12,060 |
| scCv3 | MOp | Cell | 10X v3 | 71,183 | 78 | 7,282 | 46,148 |
| snCv3Z | MOp | Nucleus | 10X v3 | 40,166 | 67 | 3,445 | 16,088 |
| AUD | AUD | Cell | 10X v2 | 71,670 | 203 | 3,969 | 10,105 |
| Isocortex-Hippocampus | 21 | Cell | SmartSeq | 827 to 16,318 | 13 to 183 | 6,099 to 9,006 | 488,099 to 2,016,775 |
| Isocortex-Hippocampus | 19 | Cell | 10X v2 | 18,307 to 216,203 | 166 to 263 | 2,874 to 4,944 | 6,102 to 15,272 |

**Figure 1.**
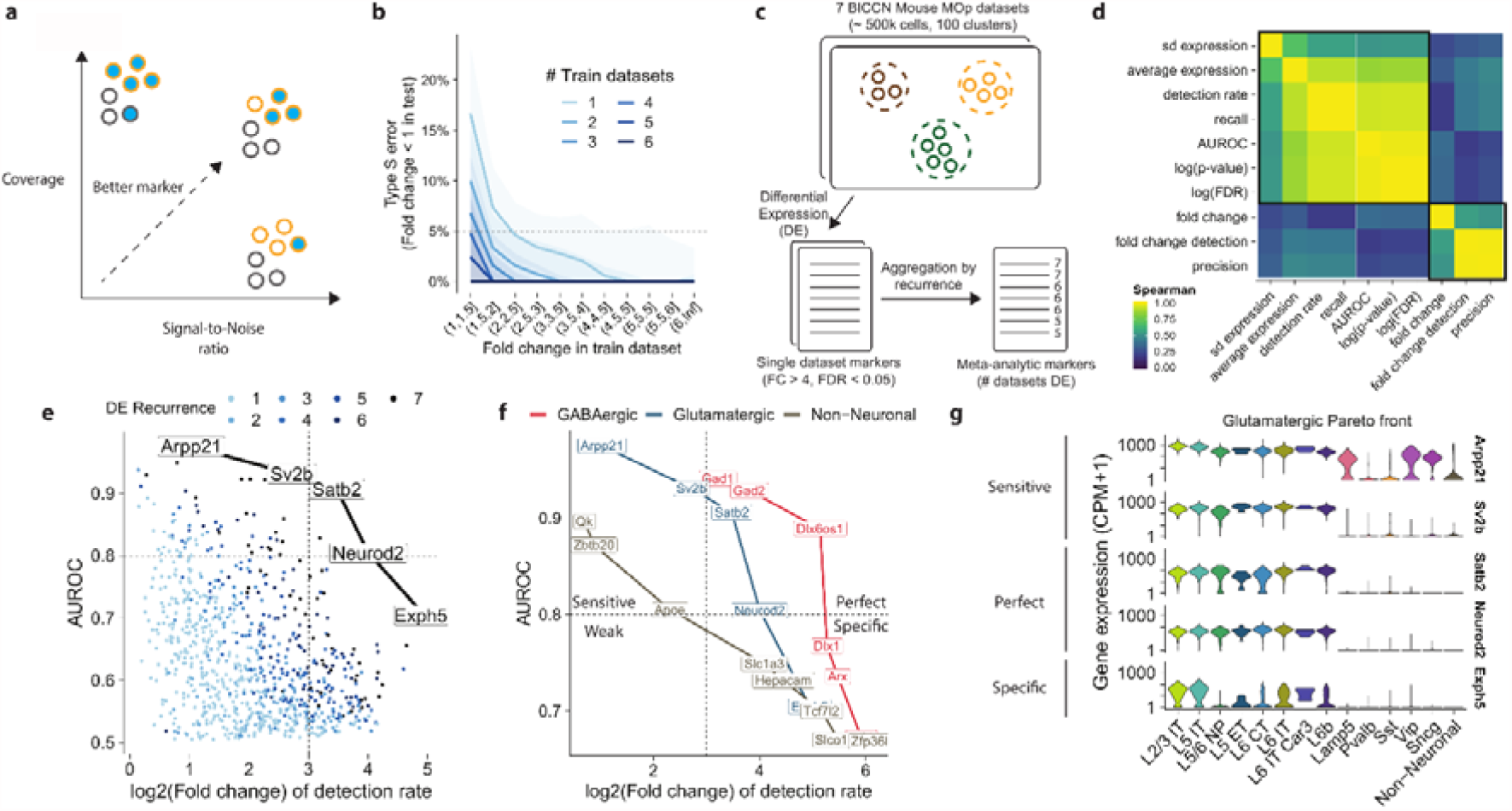
The meta-analytic Pareto front of markers: a trade-off between coverage and signal-to-noise ratio. **a** Ideal markers have high coverage (high expression in cells of interest) and high signal-to-noise ratio (relatively low expression in background cells). **b** Fraction of genes inconsistently detected as upregulated (type S error) depending on the fold change in the training dataset. Colors indicate the number of datasets used to estimate the fold change (geometric mean). **c** Schematic of extraction of meta-analytic markers: differentially expressed (DE) genes are computed independently in each dataset, meta-markers are selected based on the number of times they were DE across datasets. **d** Spearman correlation of standard DE statistics for putative markers (averaged over datasets). We highlight two independent groups of statistics that can serve as a proxy for coverage and signal-to-noise ratio. **e** Recurrent DE genes in glutamatergic neurons, using AUROC as a proxy for coverage and fold change of detection rate as a proxy for signal-to-noise ratio. Gene names and lines highlight the Pareto front of markers, which offer optimal trade-off between signal-to-noise and coverage. **f** Pareto fronts for neuronal classes (glutamatergic neurons, GABAergic neurons and non-neuronal cells) in the coverage/signal-to-noise space. We subdivide markers as perfect (high coverage and signal-to-noise), specific, sensitive, or weak (low coverage and signal-to-noise). **g** Illustration of sensitive (high target expression, some background expression), perfect (high target expression, no background expression) and specific (low target expression, no background expression) markers along the glutamatergic Pareto front.

### Meta-analytic markers are highly replicable

We started by investigating the replicability of standard differential expression (DE) statistics across BICCN datasets. Previous experiments in microarray and bulk RNAseq data by the MAQC (Shi et al. 2006) and SEQC (Consortium et al. 2014) consortia established that a fold change (FC) threshold between 2 and 4 was necessary to obtain replicable DE genes. We wondered if a similar threshold would hold for single cell RNAseq and how aggregation across datasets would improve the threshold for fold change (FC) and the area under the receiver-operator curve (AUROC), a statistic routinely used to compute the statistical significance of DE.

To assess the replicability of FC, we quantified how often one would draw inconsistent conclusions about a significant DE gene being upregulated (type S error (Gelman and Carlin 2014)). For example, given that I observed a gene with FC=2 (strongly upregulated), what is the probability that my gene will have a FC<1 (downregulated) in an independent experiment? When FC was estimated from a single dataset, as is routine in published studies, we found that a threshold of FC>4 was necessary to call a gene reliably upregulated (type S error < 5%, Fig. 1b), in line with MAQC/SEQC conclusions. In contrast, estimating FC from a higher number of datasets dramatically improved replicability: for 2 datasets the 5% error threshold is reached at FC > 2, for 3 datasets at FC > 1.5. Surprisingly, for more than 5 datasets, our results suggest that thresholding becomes unnecessary: a gene that was detected as upregulated in 5 independent datasets was almost always upregulated in the 2⍰remaining datasets, even at low effect size (FC∼1). Moreover, for a single dataset, only the top 10 upregulated genes were replicable, while the top 1000 genes are reliably upregulated when aggregating across 6 datasets (Sup. Fig. 1b). We observed similar trends for AUROCs. Based on a single dataset, the replicability threshold was AUROC>0.65, yielding 100 reliably upregulated genes. Aggregating six datasets, no replicability threshold was needed and we could identify more than 5000 reliably upregulated genes (Sup. Fig. 1a,c). The impact of dataset aggregation was particularly dramatic for small clusters and lowly expressed genes (Sup. Fig. 1d-g); for 5/85 neuron clusters, fewer than 5 of the top 10 single dataset markers (based on fold change) were reliably upregulated.

### No individual marker offers high coverage and signal-to-noise ratio

Having established that DE statistics are replicable in aggregate, we assessed a range of existing statistics and found they strongly clustered into two groups, corresponding to definitions for coverage and signal-to-noise ratio (Fig. 1c,d). The first block of statistics contained average gene expression and intuitively mapped to the notion of coverage; it also included the DE p-value and the detection rate, which are strongly indicative of genes that are broadly expressed. The second block contained the fold change and the fold change of detection rate and intuitively mapped to the notion of signal-to-noise ratio. The lack of correlation between the two blocks indicates that there is trade-off, genes have a “choice” between favoring coverage or signal-to-noise ratio. Note that this is broadly consistent with long-standing heuristic practice of considering both p-value and fold change in bulk DE through volcano plots (Cui and Churchill 2003; Goedhart and Luijsterburg 2020). In the following, we use the area under the receiver-operator characteristic curve (AUROC) as our proxy for coverage (as used in Seurat’s ROC test (Stuart et al. 2019) or LIGER’s marker detection (Welch et al. 2019; Liu et al. 2020)), fold change of the detection rate (FCd) as our proxy for signal-to-noise when we consider individual markers (as used in M3Drop (Andrews and Hemberg 2019)), and fold change (FC) as our proxy for signal-to-noise when we consider marker lists (as used in the traditional Volcano plot (Cui and Churchill 2003)).

In a FC/AUROC representation, genes offering a trade-off from best signal-to-noise marker to highest coverage marker form a Pareto front of markers (Fig. 1e). The Pareto front representation offers a rapid visualization of the strength of markers that can be associated with any given cell type. Based on our exploration of the datasets, we subdivided markers as perfect (high coverage, AUROC > 0.8, high signal-to-noise, FCd > 8), specific (high signal-to-noise), sensitive (high coverage) or weak (DE, but low coverage and low signal-to-noise). As expected, the Pareto fronts associated with Glutamatergic and GABAergic cells contain perfect markers (Fig. 1f) that identify these populations with high confidence across all technologies sampled, such as *Gad1* for GABAergic cells and *Neurod2* for Glutamatergic cells. In contrast, there is no perfect marker for non-neuronal cells: their Pareto front only includes highly sensitive markers such as *Qk* (highly expressed in non-neurons but also expressed in neurons) and highly specific markers such as the *Slco1c1* transporter (high signal-to-noise, but not covering all non-neurons), consistent with the heterogeneous nature of non-neurons and the need to use several markers in conjunction (Fig. 1f). Remarkably, the Pareto fronts were composed of perfectly recurring genes, i.e. genes that are reliably DE across all datasets (Fig. 1e, FC > 4, FDR < 0.05). Conversely, this implies that markers selected based on recurrence (number of datasets where they are reliably DE) naturally range from highly sensitive to highly specific. In contrast, high AUROC markers have high sensitivity but low specificity.

To illustrate that the chosen statistics and thresholds offer an intuitive understanding of coverage and signal-to-noise ratio, we plotted the expression of markers along the Glutamatergic Pareto front in one of the BICCN datasets (Fig. 1g). Highly sensitive markers (*Arpp21* and *Sv2b*, AUROC > 0.8) are expressed in all Glutamatergic cells at high levels but are also expressed in background cells (e.g., high expression of *Arpp21* in the *Vip, Sncg* and *Lamp5* cell types). The highly specific marker (*Exph5*, FCd > 8) is expressed almost exclusively in glutamatergic cells, but not in all cells, indicating high drop-out propensity or cell-type specific expression (e.g., it is almost not expressed in L5/6 NP). Finally, the perfect markers (*Satb2* and *Neurod2*) cover almost all cells of interest and have very limited background expression. To further investigate if our simple metrics capture known marker genes, we investigated the Pareto fronts of inhibitory subclasses as defined by the BICCN. We found that all classical markers were on the Pareto front (Sup. Fig. 1h), classified as perfect markers (*Pvalb, Lamp5*) or highly sensitive markers (Sst, Vip), with the notable exception of *Sncg*, which was only imperfectly detected in most datasets (low coverage, high signal-to-noise). A look at the Pareto front of the *Sncg* population suggests that multiple genes would offer better coverage than Sncg while preserving a high signal-to-noise ratio, in particular *Cadps2, Frem1* and *Megf10* (Sup. Fig. 1i), but that all markers tend to have some background expression in the *Vip Serpinf1* cell type. For Glutamatergic subclasses, the Pareto fronts suggested that all subclasses have perfect markers, except for IT subclasses, consistent with previous observations of gradient-like properties (Tasic et al. 2018; Yao et al. 2020a, 2020b) (Sup. Fig. 1j,k).

The FC/AUROC plot rapidly informs about the maximal strength of markers that can be expected for any given cell type. In contrast to the Volcano plot, which is based on one effect size and one significance statistic (Goedhart and Luijsterburg 2020), it relies on two effect sizes. Because we obtain replicable statistics by combining values over multiple datasets, we remove the need to visualize significance and obtain a plot with two interpretable dimensions of marker strength: signal-to-noise ratio and coverage. Typically, for each population, we suggest building the FC/AUROC plot across at least 5 datasets, identify genes on or next to the Pareto front, visualize their expression across datasets to appreciate the optimal coverage/signal-to-noise trade-off, then select the best marker(s) for the application at hand.

### The strength of individual markers decreases with finer cell type resolutions

The BICCN defined three levels of cell types: classes (such as glutamatergic neurons), subclasses (such as PV+ interneurons), and clusters (such as Chandelier cells)(Fig. 2a). While classes and subclasses had been previously experimentally characterized and showed strong statistical robustness across datasets, clusters obtained from independent datasets were more elusive (Yao et al. 2020a). To further characterize how distinct cell types are, we evaluated the number of replicable markers with increasing clustering resolution. We controlled for the increasing number of cell types by using a hierarchical approach, for example we compare a cluster to clusters from the same subclass only (Fig. 2a).

**Figure 2.**
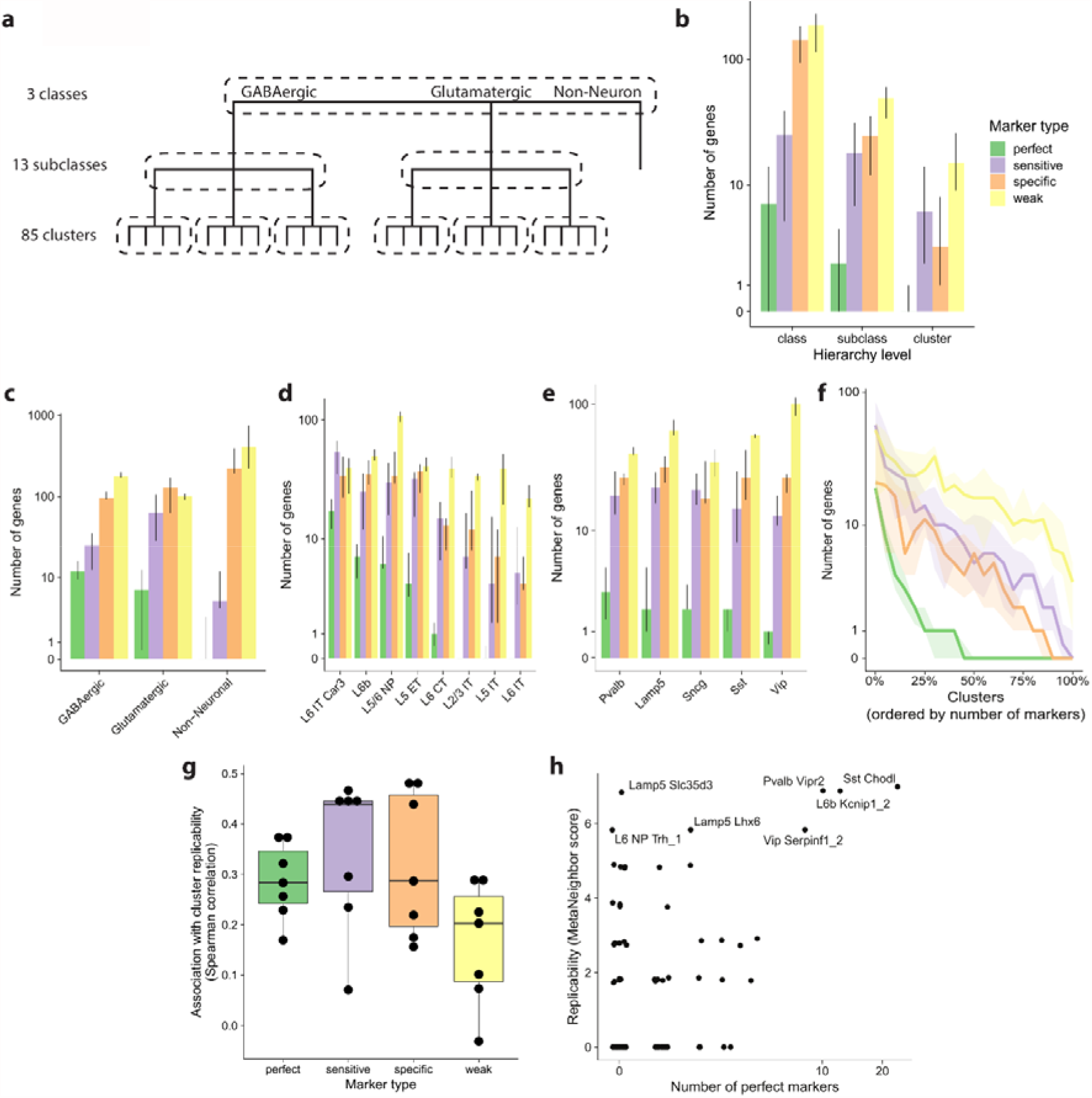
Markers are associated with higher cluster replicability, but become rare at finer resolutions. **a** Schematic of the BICCN taxonomy. Markers are selected hierarchically: each cluster is only compared to its direct neighbors in the hierarchy (dashed lines). **b** Number of reliable markers (FC>4, FDR<0.05) along the BICCN cell type hierarchy, according to marker type: perfect (AUROC > 0.8 and FCd > 8), specific (AUROC > 0.8), sensitive (FCd > 8) and weak (FDR < 0.05). **c** Number of markers of each type for BICCN classes, error bars are interquartile range across datasets. **d** Same as c for Glutamatergic subclasses. **e** Same as c for GABAergic subclasses. **f** Number of markers of each type for BICCN clusters, with cell types ordered according to number of markers. Ribbons indicate interquartile range across datasets. **g** Association between number of makers and cross-dataset MetaNeighbor replicability score at the cluster level (Spearman correlation, one dot per BICCN dataset). **h** Illustration of association of replicability (MetaNeighbor score) and number of markers in the scCv2 dataset.

To investigate how the number and quality of markers depends on the cell type hierarchy, we extracted all reliable markers (FC>4, FDR<0.05) and classified them as perfect (AUROC > 0.8 and FCd > 8), specific (AUROC > 0.8), sensitive (FCd > 8) and weak (FDR < 0.05). We observed an overall decrease in the median number of markers when going from coarse to finer resolution (397 total markers at the class level, 108 at the subclass level, 35 at the cluster level), confirming that the signal that separates neighboring populations becomes increasingly weaker (Fig. 2b). We found that all classes and subclasses had at least one perfect marker except for non-neurons and IT subclasses (Fig. 2b-e). In contrast, only around 50% of clusters had a perfect marker (Fig. 2f, Sup. Fig. 2a-d). This proportion dropped to 25% with the additional requirement that the marker should be robust across all technologies (Sup. Fig. 2a). Strikingly, a handful of clusters had extremely strong support, totaling close to 50 perfect markers in some of the datasets. Upon closer investigation, these clusters corresponded to experimentally identified populations, such as the long-projecting interneurons (Tasic et al. 2018; Paul et al. 2017)(*Sst Chodl*, up to 43 perfect markers) or Chandelier cells (Paul et al. 2017; Tasic et al. 2018)(*Pvalb Vipr2*, up to 20 perfect markers), suggesting that for these cell types, experimentally characterized differences in morphology and physiology are reflected by a high number of marker genes. Reassuringly, almost all clusters had at least one specific marker, suggesting the presence of unique characteristics (Fig. 2f, Sup. Fig. 2b).

While more data are needed to experimentally validate cell types, we wondered whether the number of markers would be predictive of computational replicability. Intuitively, a higher number of markers indicates unique aspects in a population’s transcriptional program, which should increase its identifiability across datasets. We assessed cluster replicability using MetaNeighbor, which tests the consistency of cell types across datasets using a neighbor voting framework: intuitively, if two clusters represent the same cell type, they will preferentially vote for each other (see Materials and Methods). We found that cluster replicability was indeed associated with the number of markers (rho=0.4, Fig 2g). To understand why replicability and number of markers are only partially associated, we further investigated the relationship in the best-powered datasets. We noted that, while a high number of markers was associated with higher replicability, a low number of markers did not imply low replicability (Fig 2h). Some clusters, such as *Lamp5 Slc35d3*, are found independently in all BICCN datasets and match with high statistical confidence (MetaNeighbor replicability > 0.7), despite the absence of strong markers. However, we noted that these clusters usually had a high number of specific markers (Sup. Fig. 2e). Conversely, we found some instances of clusters with low replicability and high number of markers (e.g. *Pvalb Nkx2*.*1*, Sup. Fig. 2f) but, upon further investigation, all identified “markers” were stress-related genes likely to be artefacts of the extraction protocol. Overall, the imperfect association of markers and replicability suggests that individual markers only provide a partial view of cell type identity, which is encoded broadly across the transcriptome.

### Meta-marker aggregation enables near-optimal cell type descriptions

Our previous results suggest that, at the finest level of resolution, single markers are not sufficient to unambiguously identify cell types (only ∼10 genes with AUROC > 0.8 at the subclass level, Fig. 3a). These results are consistent with the ideas that markers are affected by dropout and that clustering procedures capture information from the full transcriptome. We next tested if cell type identity can be efficiently characterized by redundant marker lists. In particular, we ask how many markers contribute to make cell types more unique, and how the selection of replicable markers improves cell type characterization.

**Figure 3.**
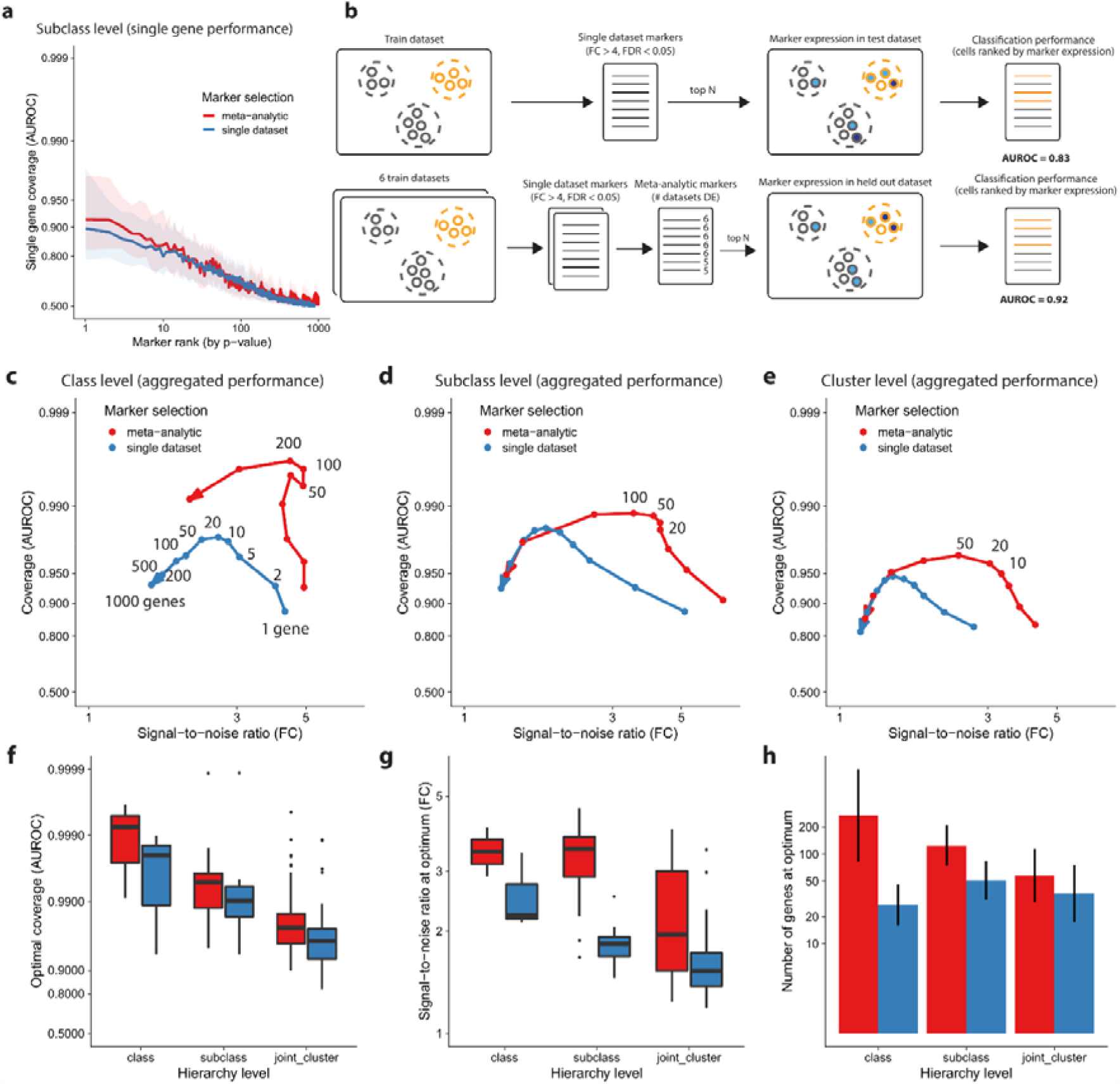
Meta-analytic aggregation of markers considerably improves the coverage/signal-to-noise trade-off. **a** Cell type classification performance of single markers as a function of marker rank. **b** Schematic of classification task using cross-dataset validation for markers from single datasets and meta-analytic markers. **c** Cell type classification performance of aggregated markers (average expression) with increasing number of markers. Performance is plotted as a parametric curve in a coverage (AUROC), signal-to-noise (fold change) space similar to Figure 1. The arrow points toward an increasing number of markers, the numbers next to the dots show the number of genes at which performance was measured (shown in full as an example for one arrow, otherwise highlighting optimal performance). **d** Same as c at the subclass level. **e** Same as b at the cluster level. **f-h** Coverage (f), signal-to-noise ratio (g) and average number of genes (h) at optimal coverage as a function of hierarchy level. Boxplots show median and interquartile range (f,g), bar plot shows mean and standard deviation (h) across datasets. In all panels, colors indicate whether markers were prioritized according to a single dataset or using the meta-analytic approach.

To study how the number of markers affects cell type identifiability, we framed gene aggregation as a classification task (Fig. 3b). How well can we predict cell type identity for the average expression of an increasing number of markers? We first created ranked marker lists for each dataset by ranking genes according to their AUROC. To test the effect of meta-analytic marker selection, we used cross-dataset validation: we computed marker replicability across 6 datasets and predicted cell types on the held-out dataset. To rank meta-analytic markers, we used two criteria: first, the number of datasets in which they were reliable DE (FC>4, FDR<0.05), second, the average AUROC. To predict cells that belong to a given cell type, we ranked cells based on the average expression of the top N markers for that cell type. We visualized performance in the FC/AUROC space, displaying classification results as a trade-off between coverage (AUROC) and signal-to-noise ratio (FC). We found that marker aggregation improved cell type identification at all levels of the hierarchy, independently of the marker prioritization strategy (Fig. 3c-e). Coverage reached an optimum between 10 and 200 genes (Fig. 3c-e), at the cost of a slightly lower signal-to-noise ratio (class, FC=6 to 6, subclass, 6 to 5, cluster, 5 to 3). Optimal performance was reached between 50 and 200 genes for classes, 20 to 100 genes for subclasses, and 10 to 50 genes for clusters.

Meta-analytic markers systematically outperformed single dataset marker genes in terms of coverage (Fig. 3f, class, AUROC=0.92 to 0.99, subclass, 0.9 to 0.99, cluster 0.85 to 0.95), signal-to-noise ratio (Fig. 3g) and number of relevant genes (Fig. 3h). In other words, the best candidates in a single dataset by a single metric are “too good to be true”. The gain in signal-to-noise ratio is particularly apparent at the cluster and subclass levels (Fig 3d-e), suggesting that the meta-analytic approach successfully extracts and combines lowly expressed markers. We checked that all results were robust to another marker prioritization strategy, where we ranked genes by fold change instead of AUROC (Sup. Fig. 3a-c).

We further investigated how the performance was distributed within hierarchy levels and across datasets (Sup. Fig. 3d-o). The overall classification performance (AUROC) increased with dataset depth (Sup. Fig. 3d). More surprisingly, the signal-to-noise ratio was approximately constant across datasets (Sup. Fig. 3h) and the number of relevant markers was slightly lower for high depth datasets (Sup. Fig. 3l). Classification performance was high for all classes and subclasses (median AUROC > 0.99, median FC > 3, Sup. Fig. 3e,f,i,j), with the notable exception of L5 IT and L6 IT (AUROC < 0.99, FC < 3). The classification performance had a wide variance at the cluster level (AUROC ranging from 0.9 to 1, FC ranging from 1.5 to 8, Sup. Fig. 3g,k), 32/85 cell types had a low signal-to-noise ratio (median FC < 2, Sup. Fig. 3g). Finally, we found that the ideal number of markers ranged from 10 to 200 and was remarkably consistent within each hierarchy level (Sup. Fig. 3l-o).

### Meta-marker enrichment is robust across datasets

Automatic annotation of cell types typically involves two steps: (a) prioritize cells that are most likely to belong to a given cell type, (b) annotate cells that exceed a pre-specified threshold condition. The threshold indicates that there is enough evidence to proceed with the annotation, for example preventing misannotation when a cell type is missing in the reference dataset. In the previous assessment, we showed that meta-analytic marker lists successfully prioritize cells, without explicit consideration for correct thresholding. We wondered whether marker expression was sufficiently consistent to be compatible with a simple thresholding method: a cell belongs to a given cell type when its marker expression exceeds the same pre-specified value for each test dataset.

For each dataset in the compendium, we computed the annotation performance at various threshold values (Fig. 4a). For example, in the *Pvalb subclass*, meta-analytic markers had a high maximal performance (F1opt>0.9) across all datasets (Fig. 4c). Additionally, the maximal performance had a distinctive plateau, indicating that a large range of thresholds had almost equivalent performance, as expected from the meta-markers’ tendency to preserve a high signal-to-noise ratio. To visualize how well optimal thresholds aligned across datasets, we defined the plateauing region as the thresholds that had at least 90% of the maximal performance (Fig. 4b). While there was a large plateau in all datasets, the plateaus did not align well, suggesting normalization issues (Fig. 4c). As a result, a meta-analytic threshold leads to good performance in most datasets, but fails in dataset with extreme properties, such as snCv2 (nuclei, 10X v2, low depth) or scCv3 (cells, 10X v3, high depth).

**Figure 4.**
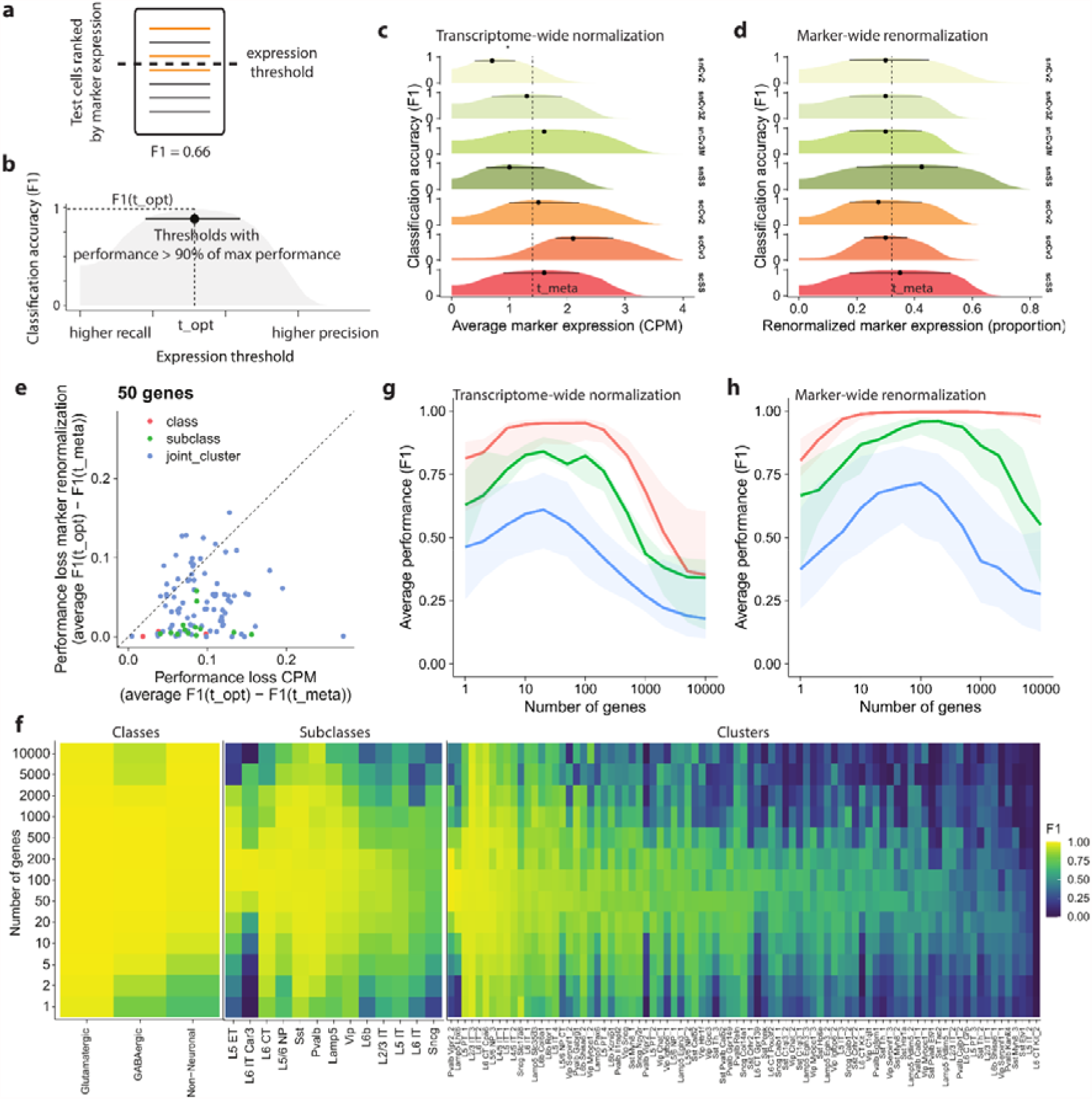
Aggregated expression of meta-analytic markers enables robust identification of cell types. **a** Schematic of threshold-based classification. The initial steps of the procedure are identical to Figure 3. **b** Illustration of statistics measuring cell type annotation performance at various thresholds. For a given dataset, cell type and number of genes (scSS dataset, Pvalb subclass, 100 genes in the illustration), F1opt is the score obtained by picking the single best threshold, indicated by a dot. The line indicates the range of near-optimal thresholds (leading to a performance higher than 0.9*F1opt). **c** Comparison of near-optimal expression thresholds across datasets (for Pvalb subclass and 100 genes). The position of the dotted line (F1meta) is obtained by averaging optimal expression thresholds across datasets. **d** Similar to b, but defining optimal thresholds based on proportion of marker expression instead of expression. **e** For each cell type, we show how much performance is lost by switching from a dataset-specific threshold (F1opt) to a single meta-analytic threshold (F1meta) for the two types of thresholds (CPM expression, marker expression proportion). Colors indicate hierarchy level, the dashed line is the identity line (performance loss is identical for the two types of thresholds). **f** For each hierarchy level, heatmap detailing classification performance for each cell type as a function of the number of genes. **g-h** Average performance as a function of the number of genes using the optimal meta-analytic expression threshold (g) or optimal marker expression proportion threshold (h). Ribbons show interquartile range across populations and test datasets.

To overcome the normalization discordance, we reasoned that the normalization issues are mainly driven by non-marker genes. Instead of considering marker expression for each cell type independently, we divided marker expression by the total marker expression (across all putative cell types). After this change, plateaus of optimal thresholds aligned across all datasets (Fig. 4d), suggesting that marker lists have preserved relative contributions in all datasets. To assess the utility of marker-wide renormalization, we directly compare how much performance is lost by switching from dataset-specific thresholds (optimal threshold per dataset) to a consensus threshold. The decrease in performance was systematically lower with marker-wide renormalization for classes and subclasses and was generally lower for clusters (Fig. 4e).

We compared the performance achieved for transcriptome-wide and marker-wide normalization as a function of the number of markers (Fig. 4g-h, Sup. Fig. 4a-b) and within each hierarchical level (Fig. 4f, Sup. Fig. 4c-e). Both methods reached high classification performance at the class and subclass level (optimal average F1 > 0.75, Fig. 4g-h), but the average performance was considerably lower at the cluster level. Marker-wide normalization yielded substantially higher classification performance (ΔF1 ∼ 0.1) and reached peak performance by successfully integrating a higher number of genes (50-500 markers, Fig. 4g-h). Performance was distributed unequally within hierarchy level, in particular for subclasses and clusters (Fig. 4f). Almost all subclasses reached optimal performance around 100-200 markers with a high performance (F1 > 0.75), with the exception of L5 IT, L6 IT and *Sncg*. At the cluster level, the performance degraded substantially: peak performance was attained around 50-100 markers, with only 43/85 of cell types reaching high performance (F1 > 0.75). All these trends were consistent with results obtained using transcriptome-wide normalization, with overall higher annotation performance (Sup. Fig. 4c-e).

### Meta-markers are enriched for genes involved in synaptic regulation and development

We next wondered if top meta-markers were enriched for specific biological processes. We performed gene set enrichment analysis for the top markers in each cell type against Gene Ontology (GO) terms from the Biological Process (BP) ontology. To focus on specific processes, we only queried terms containing between 20 and 100 genes. For each cell type, we extracted the top 3 enriched terms based on the False Discovery Rate (FDR) from the hypergeometric test. The best balance between number of enriched terms and cell type specificity was achieved for the top 100 markers for both classes and subclasses (Fig. 5a, Sup. Fig. 5a).

**Figure 5.**
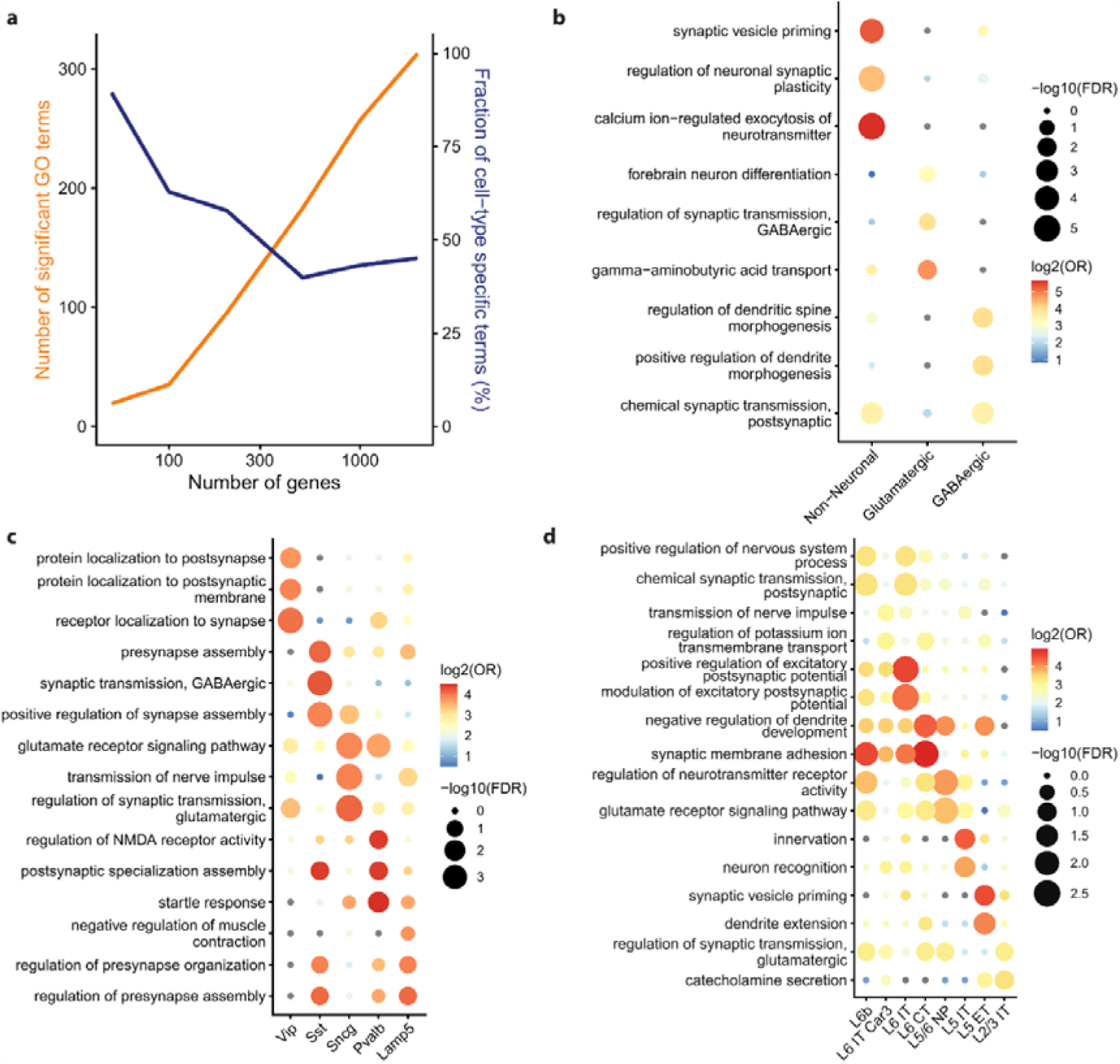
The top 100 meta-markers are enriched for specific synaptic processes. **a** Total number of significantly enriched GO terms (orange) and fraction of significant GO terms that are enriched in a unique cell type (blue) for BICCN subclasses when an increasing number of meta-markers are considered. **b** Top 3 enriched Gene Ontology (GO) terms for the top 100 meta-markers for each BICCN class. For each dot, the size reflects the False Discovery Rate (FDR), the color reflects the Odds Ratio (OR) of the enrichment test (hypergeometric test). **c** Same as b for the top 100⍰meta-markers for BICCN GABAergic subclasses. **d** Same as **b** for the top 100⍰meta-markers for BICCN Glutamatergic subclasses (only top 2 terms per cell type are shown).

We found strong enrichment for synaptic properties for all cell types. At the class level, non-neuronal markers were enriched for synaptic support functions, such as “Regulation of neuronal synaptic plasticity” (Fig. 5b). Glutamatergic neurons were enriched for synaptic regulation (such as the regulation of GABAergic transport), while GABAergic neurons were enriched for gene sets related to the regulation of spine and dendrite formation. At the subclass level (Fig. 5c,d), GABAergic neurons were most distinguishable based on synaptic sub-properties, such as localization to synapse (*Vip*), synapse assembly (*Sst, Lamp5*), or glutamate transmission regulation (*Sncg*). Glutamatergic subclasses showed a similar enrichment of synaptic sub-properties, including various aspects of potential regulation and synaptic transmission (L6b, L6 IT, L2/3⍰IT, L5/6 NP), as well as synaptic development (L6 CT, L5 IT, L5 ET). We further confirmed that these findings were consistent with the enrichment of the top 200 markers, which also highlighted gene sets involved in synaptic regulation and development (Sup. Fig. 5b-d). These results suggest that meta-markers define a plausible biological subspace revealing cell type differences in terms of synaptic properties.

### Meta-markers improve deconvolution performance

One of the primary purposes to which cell atlas data may eventually be put is deconvolution of bulk data where cell composition is likely related to the condition of interest (e.g., disease). Single cell data have been routinely used to increase deconvolution performance in recently developed tools (Tsoucas et al. 2019; Wang et al. 2019; Newman et al. 2019; Dong et al.), but performance remains plagued by batch effects and cell type similarity (Newman et al. 2019; Huang et al. 2020; Cobos et al. 2020). The role of marker genes in deconvolution remains particularly unclear: a recent benchmark suggests that the quality of markers is more important than the deconvolution method (Cobos et al. 2020), in most studies the influence of the number of markers is only partially assessed (Newman et al. 2019; Hunt and Gagnon-Bartsch 2019). Our annotation assessment suggested that cell types are best captured with 10 to 200 meta-analytic markers; deconvolution is a natural place to test this heuristic.

To measure the number of genes that yield maximal deconvolution performance, we generated thousands of pseudo-bulk datasets with known mixing proportions from each of the BICCN datasets (Sup. Fig. 6a). As in previous experiments, we directly compared the performance of markers extracted from single datasets and performance of meta-analytic markers. We initially compared two tasks: (a) within-dataset cross-validation, where cell type profiles are learned from a training fold and tested on a held-out set from the same dataset, (b) cross-dataset-validation, where profiles are learned on one dataset and tested in another dataset. Within-dataset cross-validation proved to be a simple task, yielding extremely high performance (Pearson ∼ 1, not shown). In contrast, cross-dataset-validation showed only modest performance (Pearson ranging from 0 to 1, Sup. Fig. 6b), highlighting the difficulty of the deconvolution task. Because deconvolution applications typically involve different datasets, we focused our analyses on cross-dataset validation.

**Figure 6.**
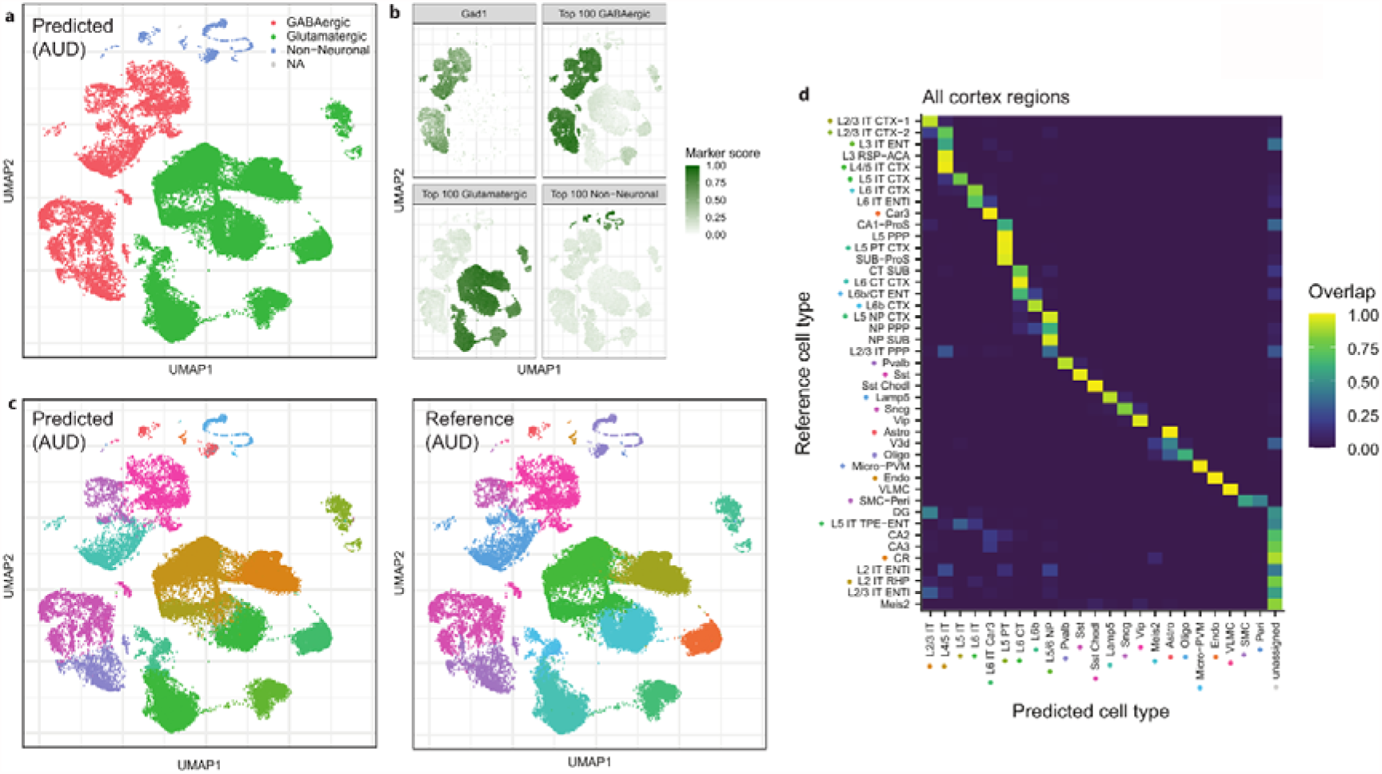
Meta-analytic markers from the primary motor cortex (MOp) generalize to other cortical regions. **a** Example of class-level predictions in the auditory cortex (AUD), where cells are embedded in UMAP space and colored according to predictions based on the top 100 meta-markers for MOp classes. Cells are assessed independently and remain unassigned (NA) if the marker enrichment score is lower than 1.5 for all classes. **b** Marker scores (re-normalized between 0 and 1) used to determine cell type assignments. The first subpanel shows the score obtained from a single GABAergic marker, the three other panels show the scores obtained by combining the top 100 meta-markers for each class. **c** Subclass-level predictions in the auditory cortex based on the top 100 MOp meta-markers (left) and reference labels (right). Cells remain unassigned (NA) if the marker enrichment score is lower than 1.5 for all subclasses. See panel **d** for color legend (some reference cell types are absent from AUD). **d** Confusion matrix showing the concordance of subclass-level predictions based on the top 100 meta-markers with reference cell types across 40 brain areas. Cells are unassigned if the marker enrichment is lower than 1.5 for all subclasses.

Deconvolution performance rapidly degraded along the neuron hierarchy (Sup. Fig. 6b), ranging from almost perfect performance for classes (Pearson ∼ 1) to low performance for clusters (Pearson < 0.5). Classes were easily learnable across all tasks (Sup. Fig. 6c), even using random genes, suggesting that at this level of the hierarchy, cell types are strongly uncorrelated and can easily be separated along the first principal component. At the subclass level, the performance of random genes remained close to 0, suggesting stronger covariation compared to classes (Sup. Fig. 6d). Meta-analytic markers yielded more robust deconvolution performance, with performance increasing up to 100 genes, while markers from single datasets prioritized only around 10 informative genes. The trend was similar at the cluster level, but with lower overall performance (Sup. Fig. 6e). Meta-analytic markers again proved to be more robust, prioritizing around 50 informative genes compared to 10-20 from single datasets. Overall, our results suggest that, in conjunction with batch effects, the increasing covariation of cell types at finer resolution significantly complicates the deconvolution task. The prioritization of a large number of robust marker genes is an important first step towards successful deconvolution.

### Meta-markers reveal a generalizable description of cell types

We have previously shown that meta-marker signatures generalize across laboratories and technologies. We next asked how well they generalize across the cortex by predicting cell types in a BICCN dataset combining multiple cortical and hippocampal brain regions (Yao et al. 2020b). To understand how easily meta-markers generalize, we used a straightforward annotation method: assign cells to the cell type with the highest average meta-marker expression. For simplicity, we considered the same number of meta-markers for all cell types: 100 at the class and subclass level, 50 at the cluster level.

We started by predicting cell types in the auditory cortex sub-dataset, containing 71,670 cells annotated to 203 cell types. We chose to focus on the auditory cortex because of its large number of cells, and to investigate the generalizability of cell types derived from a motor area (MOp) to a sensory area. While inhibitory cell types are expected to generalize, excitatory cell types have been shown to have divergent patterns (Tasic et al. 2018). At the class level, top meta-markers enabled perfect classification down to every single cell (Fig. 6a). Assignments can be traced back to meta-marker scores, as well as individual genes (Fig. 6b). Consistent with our previous points, the GABAergic score is uniformly high across all cells labeled as GABAergic. In contrast, the expression of the single best marker, *Gad1*, is more variable in GABAergic cells and displays sporadic expression in non-GABAergic cells.

Remarkably, at the subclass level, meta-markers enabled similarly strong cell type assignments, as suggested by the uniform coloring of clusters in UMAP space (Fig. 6c). Note that the assignments occur in each cell independently, without knowledge about clusters or expression profiles of neighboring cells, highlighting the consistency of meta-marker expression. This procedure allowed the identification of rare cell types, even when only one or two cells were present in the dataset (e.g. smooth muscle cells and pericytes, Sup. Fig. 7b). The predicted assignments corresponded almost perfectly to the reference cell types (Fig. 6c). The main exception were deep layers IT cell types, in particular one group of L5 IT cells tended to be assigned as L4/5 IT or L6 IT (Sup. Fig. 7c). Finally, cluster-level predictions also proved extraordinarily consistent, with smooth transitions between cell types that mapped with auditory cortex reference annotations (Sup. Fig. 8).

To further highlight high-quality predictions, we quantified assignment confidence using meta-marker enrichment (observed expression over expected expression) as a “QC” metric. In the auditory cortex, we found that a threshold of 1.5 offered an optimal trade-off between annotation recall and precision (Sup. Fig. 9a-c). Raising the threshold to 2 further selected high-confidence calls, yielding higher precision for slightly lower recall. Interestingly, cells that became unassigned were mostly located in regions where predictions and reference disagreed: deep IT layers, and inhibitory neurons bridging medial ganglionic eminence (MGE) and caudal ganglionic eminence (CGE) subclasses (Sup. Fig. 7a). Meta-marker enrichment thus offers a good proxy for prediction quality, enabling to identify cells with a high-confidence cell type assignment.

Next, we systematically quantified the agreement of meta-marker based predictions and reference annotations across all brain regions and 43 consensus subclasses. We found exceptionally good agreement, with all reference subclasses mapping to exactly one predicted MOp subclass (Fig. 6d). All MOp subclasses matched strongly with their “natural” counterparts in the reference dataset, such as “L2/3 IT” with “L2/3 IT CTX-1”. Remarkably, reference cell types absent in MOp (such as hippocampal cell types) were mostly labeled as “unassigned”, suggesting that meta-marker signatures correctly avoid labeling unseen cell types. This trend became particularly obvious for cells with marker enrichment > 2 (Sup. Fig. 9d), where all unseen cell types became “unassigned”, while conserving high matching scores between shared cell types.

## Discussion

By assessing marker replicability across 7 datasets, we selected robust markers and identified the optimal number of markers to define a cell type. We identified highly replicable markers for 85 cell types from the mouse primary motor cortex (Sup. Data 1-3). This meta-analytic strategy proved particularly important for rare populations and lowly expressing genes (Fig. 1). Compared to previous efforts (Tasic et al. 2018; Mancarci et al. 2017; Yao et al. 2020a), we identified a high number of robust markers at high cell type resolution: at the BICCN cluster level, cell types were best characterized by 10-200 meta-analytic markers, a two-fold increase of reliable markers compared to markers selected from single datasets (Fig. 3). Interestingly, we found that only 50% of clusters had strong markers (Fig. 2), but that some of the clusters lacking strong markers (e.g. *Lamp5* Slc35d3) were consistently identified in all BICCN datasets, suggesting broad encoding of their identity and highlighting the need of extended marker signatures.

We found that the simple aggregation of marker expression enabled the annotation of individual cells (Fig. 4), suggesting that careful feature selection is enough to provide a rough definition of cell types. Remarkably, marker lists derived from a single cortical region generalized with high accuracy to other cortical regions without any methodological fine-tuning (Fig. 5). By introducing redundant information about cell types, meta-analytic markers dramatically increased cell type separability (Fig. 3). However, adding more markers is only beneficial if they are cell type-specific. As a result, we established that the ideal number of markers decreases with cell type resolution: 200 genes to separate classes (lowest resolution, e.g. GABAergic neurons), 100 genes for subclasses (e.g., *Pvalb* interneurons) and 50 genes for clusters (highest resolution, e.g., Chandelier cells).

By combining datasets that were generated using different technologies, the markers we propose are likely to generalize well with respect to this axis of variation. Moreover, we show that our marker descriptions generalize to other cortical regions, despite all “training” datasets sampling from the same cortical region. However, the data used in this study were obtained from adult mice with limited genetic background and grown in lab conditions. As a result, it remains unclear how well the marker descriptions would generalize across development or biological conditions. On the other hand, as our approach relies on a simple procedure, marker lists can easily be extended to incorporate new sources of variation, such as additional brain regions, species or biological conditions. On a similar note, markers depend on one particular annotation effort, but we can expect the neuron taxonomy to evolve with additional data, in particular the fine-resolution clusters. Our framework, available as an R package, allows to rapidly evaluate the consistency of marker expression for new cell type annotations.

To highlight the replicability of marker descriptions, the manuscript relies on simple methods, but marker lists can easily be combined with more sophisticated methods for marker selection or cell type assignment. For example, experimental applications routinely require either a few specific markers to target one cell type (Huang 2014) or a panel of hundreds of markers to jointly separate all cell types (Moffitt et al. 2018). Marker lists can be combined with methods to select concise sets of markers (Asp et al. 2019; Zhang et al. 2020; Dumitrascu et al. 2019) by filtering candidates that are likely to generalize. Similarly, development studies (Hobert 2008; Huang 2014; Kessaris et al. 2014; Lodato and Arlotta 2015; Mayer et al. 2018; Tosches et al. 2018) indicate that neural lineages are marked by the specific onset and offset of key transcription factors (TFs), but the expression of these key TFs may not be maintained at later stages or only at low levels. Since our approach is powered to identify lowly expressed markers, it can be combined with time series data to help identify replicable lineage-specific genes.

This study focused on the neuron hierarchy, but our strategy generalizes to other tissues. In order to encourage broader adoption, we have made our code available as a package and in the vignette we show how our analyses and guidelines can be similarly applied to a pancreas compendium. We chose to focus on the BICCN dataset because of its complexity (85 neuronal cell types), comprehensiveness (∼500,000 cells with latest sequencing technologies) and diversity (6 technologies used). Our results suggest that, in the brain, there is a clear separation at the top two levels of the hierarchy (3 classes, 13 subclasses), but that the molecular signature of half the clusters remains unclear. We expect that similar conclusions can be drawn for other tissues, such as blood, where there is a similar hierarchical organization of cell types. The main difficulty is to identify replicable cell types across datasets, which may be challenging during development or complex differentiation processes, such as hematopoiesis.

The selection of replicable markers from single cell atlases is a promising avenue for several applications, including cell type annotation, selection of gene panels and bulk data deconvolution. It reduces rich information to a prioritized list that is simple to use and to refine. New computational methods will benefit from highly condensed prior information about genes in the cell type space, without having to train on large reference datasets. Finally, as new datasets are generated, marker lists will become increasingly robust to new sources of variation, leading to higher downstream performance across a diverse array of tasks.

## Materials and Methods

### Datasets

We downloaded the mouse primary cortex (MOp) BICCN datasets and cell type annotations from the NeMO archive (http://data.nemoarchive.org) according to preprint instructions (Yao et al. 2020a). We considered the 7 transcriptomic datasets from the mouse primary cortex: single cell Smart-Seq (scSS), single nucleus Smart-Seq (snSS), single cell Chromium v2 (scCv2), single nucleus Chromium v2 (snCv2), single cell Chromium v3 (scCv3), single nucleus Chromium v3 from the Macosko and Zeng labs (scCv3M and scCv3Z, respectively)(Table 1). We kept all cells with “class” annotated as “Glutamatergic”, “GABAergic” or “Non-Neuronal” and kept genes that were common to all datasets, arriving at a total of 482,712 cells and 24,140 genes. We normalized counts to counts per millions (CPM). For cell types, we considered five levels of annotations provided by the BICCN: “class”, “subclass”, “cluster”, “joint_subclass” and “joint_cluster”. “subclass” and “cluster” labels were obtained by clustering and annotating the datasets independently, while “joint_subclass” and “joint_cluster” labels were obtained through joint clustering and annotation. Throughout the manuscript, we use “joint_cluster” labels when we need common annotations across datasets, otherwise, we use “cluster” labels. To map “subclass” labels across datasets, we used the independent clustering, but mapped all clusters to one of the following names: “L2/3 IT”, “L5 ET”, “L5 IT”, “L5/6 NP”, “L6 CT”, “L6 IT”, “L6 IT Car3”, “L6b”, “*Lamp5*”, “*Pvalb*”, “*Sncg*”, “*Sst*”, “*Vip*”. In the last section (generalizability of meta-markers), we use the “joint_subclass” annotation instead, because it explicitly includes the distinction between L4/5 IT and L5 IT cells.

The BICCN isocortex and hippocampus dataset was downloaded from the NeMO archive (http://data.nemoarchive.org)(Yao et al. 2020b). The full dataset contains 1,646,439 cells annotated to 379 cell types. Due its size, it was separated into sub-datasets corresponding to individually sequenced brain regions (as annotated in the “region_label” metadata column), resulting in 19 brain regions sequenced with 10X v3, 21 brain regions sequenced with SmartSeq (Table 1). We subset all datasets to a common set of 24,140 genes. Preprocessing was similar to the MOp datasets: we kept all cells with “class” annotated as “Glutamatergic”, “GABAergic” or “Non-Neuronal” and normalized counts to counts per million (CPM) for SmartSeq datasets or counts per 10,000 (CP10K) for 10X datasets.

### Meta-analytic hierarchical differential expression statistics

For each cell type, we computed DE statistics independently in each dataset using MetaMarkers’ “compute_markers” function. We compared a cell type to neighboring cell types in the BICCN taxonomy by setting the “group_labels” parameter. For example, the “GABAergic” class contains the “*Pvalb*”, “*Sst*”, “*Sncg*”, “*Lamp5*” and “*Vip*” subclasses. By stratifying analysis by classes, DE statistics for “*Pvalb*” were obtained by comparing “Pvalb” cells to all cells that are either “*Sst*”, “*Sncg*”, “*Lamp5*” or “*Vip*”, but ignoring cells from other classes (excitatory neurons and glia). At the cluster level, analysis is stratified by subclasses, e.g., *Pvalb* subtypes are compared to other *Pvalb* subtypes only.

For each dataset, “compute_markers” returns a table of standard statistics. Let *x*_*ij*_ be the expression of gene *i* in cell *j* (normalized to CPM in all the manuscript), let *C* be the cells belonging to the cell type of interest, and 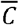 be all background cells. All statistics are computed for each gene independently, so we will drop the subscript *i* in the following. The fold change (FC) is computed as the ratio of average expression between the cell type of interest and background cells, 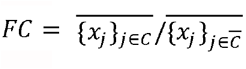. Statistical significance is based on the ROC test. First we compute the AUROC according to the following formula (derived from the Mann-Whitney U statistic):

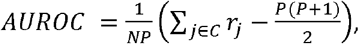

Where *P* =|*C*| are the number of positives (cells from the cell type of interest), 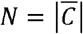 are the number of negatives (background cells), and *r*_*j*_ are the ranks of positives (obtained after ranking all cells according to the gene’s expression value). P-values are computed under a normal approximation of the AUROC with continuity and tie correction according to the following formulas:

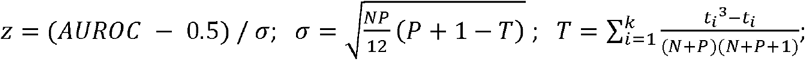

where *z* follows a standard normal distribution under the null hypothesis that positives and negatives are from the same population, *σ* is the analytical standard deviation of AUROC, *T* is a tie correction formula where *k* is the number of distinct expression values and *t*_*i*_ is the number of cells that share the same expression value with index *i*. P-values are converted to False Discovery Rates (FDR) according to the Benjamini-Hochberg procedure. For exhaustivity, we considered four additional statistics related to binarized gene expression: gene detection rate, fold change of detection rate (FCd), recall and precision. Gene detection rate is the fraction of cells in the population of interest that express the gene of interest, 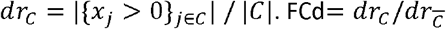 is the ratio of gene detection rates in the population of interest over the background population. Recall is identical to gene detection rate (seen from a classification perspective). Precision 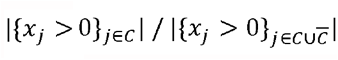 is the fraction of cells expressing the gene of interest that belong to the population of interest. All operations are vectorized across genes and cell types to allow rapid marker extraction and aggregation across datasets.

We combined statistics across datasets using MetaMarkers’ “make_meta_markers” function, which averages the above statistics across datasets for all cell types. “make_meta_markers” uses the arithmetic mean by default, and uses the geometric mean for the following statistics: FC, FCd, expression. To define DE recurrence, we used the number of datasets where a gene is reliably DE (“fdr_threshold=0.05”, “fc_threshold=4”). Throughout the manuscript, we considered a gene to be DE if it had a FC>1 and an FDR<0.05, and reliably DE if FC>4 and FDR<0.05.

### Reliable fold change and AUROC thresholds

To establish the reliability of FC, we picked all combinations of training datasets and extracted genes that were significantly upregulated in all training datasets (AUROC>0.5, FDR<0.05, average FC>1). Then, for each gene, we looked up the held out datasets and counted how often the gene remained upregulated (FC>1) or was detected as downregulated (FC<=1). We summarized the results as a type S error, the fraction of held out datasets where the gene was detected as downregulated. Formally, let *G* be the set of genes that are consistently upregulated across training datasets *d*_1_, …, *d*_*m*_. Let *d* ′ _1_, …, d′ _*n*_be the held-out test datasets. For a given cell type, the average type S error is defined as:

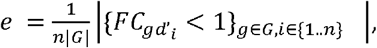

where 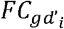 is the fold change of gene *g* in test dataset *d′*_*i*_. We computed the type S error across all combinations of cell types and training datasets. To establish the reliability of AUROC, we followed a similar procedure, replacing the FC<1 condition by AUROC>0.5.

### MetaNeighbor cell type replicability score

To compute the association between the number of markers and cell type replicability, we computed cell type similarity using MetaNeighbor by following the procedure described in (Yao et al. 2020a). Briefly, MetaNeighbor uses a neighbor voting framework to match cell types from a train dataset to a test dataset, where the matching strength is quantified as an AUROC. First, we use the “MetaNeighborUS” function to create a graph where each node is a cell type and each edge is the matching strength (directed from train dataset to test dataset). By applying the “extractMetaClusters” function, we keep only edges that correspond to high confidence reciprocal matches (1-vs-best AUROC > 0.7 both ways). After this step, we are left with groups of connected cell types that we call “meta-clusters”. The replicability score is the number of datasets spanned by the meta-cluster, e.g. a cell type has a score of 6 if it is connected to cell types from 5 other datasets. For visualization purposes, we created jittering by adding the average AUROC across the meta-cluster to the replicability score. To avoid overfitting, we considered the “cluster” annotations from the BICCN, which were obtained by clustering and annotating the datasets independently.

### Marker-based cell type classification

To quantify the ability of a list of markers to identify a cell type, we framed the problem as a hierarchical classification task where we predict cell type labels from gene expression. First, for each cell, we computed a prediction score by averaging expression profiles across markers. Let *x*_*ij*_ be the CPM-normalized expression of gene *i* in cell *j*, and *M*_*c*_ be a set of marker genes for cell type *c*. For each cell *j*, we compute the marker score is:

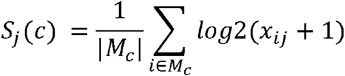

This score is efficiently implemented by MetaMarker’s “score_cells” function. To obtain marker-wide renormalized scores, we compute the above score for a series of cell types *c*_1_,..,*c*_*n*_ then, for each cell type, we compute:

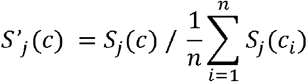

To compute classification performance, we labeled cells from the cell type of interest as positives and cells from cell types sharing the same parent class or subclass as negatives (similar to DE statistics computation, see “Meta-analytic hierarchical differential expression statistics”). Intuitively, we are looking whether positives (cells from the cell type of interest) have high prediction scores (marker scores). We summarized the prediction accuracy as an AUROC (in the threshold-free case) and F1 (harmonic mean of precision and recall, in the thresholding case). To avoid circularity, we always made predictions on held out datasets. For markers from a single dataset, predictions were averaged across the 6 remaining datasets; for meta-analytic markers, we picked markers on all combinations of 6 datasets and predicted cell types in the remaining dataset. We obtained classification scores for individual populations of neurons by averaging over every combination of train and test datasets.

### Gene ontology enrichment of meta-markers

Gene ontology terms and mouse annotations were downloaded using the org.Mm.eg.db and GO.db R packages. To focus on specific cell processes, we further selected terms from the “Biological Process” ontology containing between 20⍰and 100 gene annotations. Gene set enrichment was computed using the hypergeometric test, based on R’s “phyper” function and the Maximum Likelihood Estimate (MLE) of the sample odds ratio (OR).

### Marker-based deconvolution

To investigate the impact of marker selection on deconvolution, we applied deconvolution in a hierarchical framework similar to DE computation and cell type classification. We applied Non-Negative Least Square (NNLS) deconvolution (Abbas et al. 2009) using the nnls R package, which was shown to be both efficient and accurate according to multiple recent benchmarks (Patrick et al. 2020; Cobos et al. 2020). Briefly, we inferred cell type proportions from the following equation:

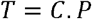

where T is a bulk expression matrix (genes x sample), in our case pseudo-bulk matrices extracted from each test dataset, C is a cell type signature matrix (genes x cell type), P is the estimated cell type proportion matrix (cell type x sample). To test all combinations of train and test datasets, we split each dataset in half by assigning each cell randomly to a test or train fold. From each train fold, we built signature matrices by averaging unnormalized expression profiles for each cell type. From each test fold, we built 1000 pseudo-bulks containing 1000 cells. To generate pseudo-bulks with highly variable cell type proportions, we started by drawing target cell type proportions for each pseudo-bulk, in a procedure similar to (Cobos et al. 2020). We sampled target proportions for each cell type from a uniform distribution, normalized proportions to 1, then converted to a target number of cells which we sampled with replacement, then averaged the unnormalized counts.

Given a set of markers (obtained from a single dataset or meta-analytically across all datasets except the test dataset), a train dataset (signature matrix) and a test dataset (1000 pseudo-bulks), we performed NNLS deconvolution by subsetting the signature matrix C and pseudo-bulks T to the set of markers. We computed deconvolution performance as the Pearson correlation between theoretical cell type proportions and the predicted cell type proportions (one value per pseudo-bulk). For computational efficiency, we only tested one group of populations at the “joint cluster” level. We chose to focus on the Lamp5 populations, as it contained 8 populations that were well represented across all datasets (range 14 to 3016 cells per single population, 257 cells on average).

Note that, because of the difficulty of matching UMI counts with full-length read counts (Newman et al. 2019), we only considered train-test combinations within similar technologies (one pair of Smart-seq datasets, 10 pairs of 10X datasets). To control for globally encoded differences in expression profiles (correlating with the first principal component), we created random marker sets by picking genes that were expression-matched with meta-analytic markers (decile-matched).

### Generation of robust meta-marker sets

We generated meta-marker sets for each cell type in the MOp hierarchy (Sup. Data 1-3), using the “class”, “joint_subclass” and “joint_cluster” annotation levels (see “Meta-analytic hierarchical differential expression statistics”). We kept meta-markers that were either strongly DE (FC>4, FDR<0.05) in at least one dataset or had a meta-analytic FC > 2. We ranked the remaining markers by recurrence, then by AUROC, and selected the top 100 ⍰genes (top 50 genes for clusters). If fewer than 100Cmarkers remained, we selected all remaining markers.

### Cell type annotation of the BICCN isocortex and hippocampus datasets

We annotated cells in the isocortex and hippocampus datasets using our robust marker lists (see “Generation of robust marker lists for all BICCN MOp cell types”). To annotate cell types, we adopted a hierarchical cell type annotation procedure. We classified each brain region independently, starting from the log-normalized count matrix. First, we obtained marker scores (average meta-marker expression, see “Marker-based cell type classification”) for all cells by running MetaMarker’s “score_cells” function. Then, marker scores were converted into cell type predictions using MetaMarker’s “assign_cell” function, which finds the marker set with the highest score and returns several QC metric, including the highest score and the marker enrichment (observed score divided by expected score, under the assumption that all marker sets have equal expression). The “assign_cell” function takes two parameters: marker scores and group-level assignments. For subclasses, we provided class-level predictions as the group assignments; for clusters, we provided subclass-level predictions as the group assignments. To filter out cells with unclear assignments, we labeled cells that had a marker enrichment below 1.5 (unless otherwise indicated in the text) as “unassigned”.

## Data and code availability

The datasets analyzed during the current study are available in the NeMO archive (https://nemoarchive.org/) at https://assets.nemoarchive.org/dat-ch1nqb7. The full meta-marker lists for the BICCN cell types and optimal number of markers are available on FigShare at https://doi.org/10.6084/m9.figshare.13348064. The code for MetaMarkers is freely available as an R package on Github at https://github.com/gillislab/MetaMarkers.

## Acknowledgments

JG was supported by NIH grants R01MH113005 and R01LM012736. SF was supported by NIH grant U19MH114821.

## Author contributions

SF and JG designed the experiments, performed the data analysis and wrote the paper. All authors read and approved the final manuscript.

## Supplemental Material

**Supplemental Figure 1. a-c** Type S error as a function of AUROC in train datasets (a), marker rank by fold change (b) and marker rank by AUROC (c). The dashed line indicates a type S error of 5%, ribbons around lines indicate variability across cell types and test datasets. **d-g** Type S error as a function of AUROC (d) or FC (e-g) in train dataset, with facets showing variability across hierarchy level (d,e), average cell type size (f) and average gene expression (g). **h** Pareto fronts in FC/AUROC space for inhibitory subclasses. Arrows point to the main historical marker for each subclass. **i** Expression of genes on the Sncg Pareto front across BICCN inhibitory clusters. **j** Pareto fronts in FC/AUROC space for excitatory subclasses. **k** Expression of genes on the L5 ET Pareto front across BICCN excitatory clusters.

**Supplemental Figure 2. a-d** Number of perfect markers (a), specific markers (b), sensitive markers (c), and weak markers (d) for BICCN clusters, with cell types ordered according to number of markers, colored according to the dataset used to compute markers. **e-f** MetaNeighbor replicability as a function of the number of specific markers in the scCv2 dataset (e) and the number of perfect markers in the snCv3M dataset (f).

**Supplemental Figure 3. a-c** Parametric curve in FC/AUROC space showing evolution of classification performance with an increasing number of marker genes at the class (a), subclass (b) and joint cluster (c) level. **d-g** Breakdown of optimal AUROC performance (meta-analytic markers) as a function of dataset depth, colored by hierarchy level (d), for individual classes, showing variability across test datasets (e), for individual subclasses, showing variability across test datasets (f), for individual clusters, showing variability across test datasets (g). **h-k** Same as d-g with signal-to-noise ratio (FC) at optimal performance instead of AUROC. **l-o** Same as d-g with number of genes at optimal performance instead of AUROC.

**Supplemental Figure 4. a-b** Summary of optimal classification performance (F1) across hierarchy levels with transcriptome-wide normalization (a) and marker-wide renormalization (b). Variability is shown across cell types and test datasets. **c-e** Heatmap detailing classification performance for each cell type as a function of the number of genes at the class (c), subclass (d) and cluster (e) level.

**Supplemental Figure 5. Top 200 meta-markers show strong, but less specific, enrichment for synaptic processes. a** Total number of significantly enriched GO terms (orange) and fraction of significant GO terms that are enriched in a unique cell type (blue) for BICCN classes when an increasing number of meta-markers are considered. **b** Top 3 enriched Gene Ontology (GO) terms for the top 200 meta-markers for each BICCN class. For each dot, the size reflects the False Discovery Rate (FDR), the color reflects the Odds Ratio (OR) of the enrichment test (hypergeometric test). **c** Same as **b** for the top 200⍰meta-markers for BICCN GABAergic subclasses. **d** Same as **b** for the top 200⍰meta-markers for BICCN Glutamatergic subclasses (only top 2 terms are shown).

**Supplemental Figure 6. Meta-analytic markers improve deconvolution performance at every level of the hierarchy. a** Schematic of deconvolution task. **b** Summary of deconvolution performance (Pearson’s r) at each hierarchy level with 100 markers per cell type. Colors show 3 marker prioritization strategies (single dataset markers, meta-analytic markers or expression-level matched random genes). **c-e** Deconvolution performance (Pearson correlation of true and estimated cell type proportions) for 3 marker prioritization strategies at the class level (c), the subclass level (d), and the cluster level (e). Colors as b.

**Supplemental Figure 7. Focus on subclass-level predictions in the auditory cortex**. a Subclass-level predictions in the auditory cortex based on the top 100 meta-markers. Cells remain unassigned (NA) if the enrichment score is lower than 2 for all subclasses. **b** Subclass-level predictions for non-neurons in the auditory cortex based on the top 100 meta-markers (left) and reference labels (right). **c** Subclass-level predictions for Intra-Telencephalic (IT) excitatory neurons in the auditory cortex based on the top 100 meta-markers (left) and reference labels (right).

**Supplemental Figure 8. Cluster-level predictions in the auditory cortex. a-c** Cluster-level predictions for Lamp5 inhibitory neurons (a), Near-Projecting (NP) excitatory neurons (b) and layer 2/3 Intra-Telencephalic (IT) excitatory neurons (c) in the auditory cortex based on the top 100 meta-markers (left) and reference labels (right). In all panels, cells remain unassigned (NA) if the enrichment score is lower than 1.5 for all clusters.

**Supplemental Figure 9. The marker enrichment score provides robust separability of cell types in other cortical regions. a** Marker enrichment scores based on the top 100⍰meta-markers for the 3⍰BICCN classes in the auditory cortex. The facets are organized according to reference cell types (from the auditory cortex), the x-axis according to meta-markers sets (for the motor cortex). **b** Same as a for BICCN inhbitory subclasses. **c** Same as **a** for BICCN excitatory subclasses. **d** Confusion matrix showing the concordance of subclass-level predictions based on the top 100 meta-markers with reference cell types across 40 brain areas. Cells are unassigned if the marker enrichment is lower than 2 for all subclasses.

**Supplemental Data 1. Class-level markers**. Top 100 robust markers for BICCN cell types at the class level in CSV format (.csv).

**Supplemental Data 2. Subclass-level markers**. Top 100 robust markers for BICCN cell types at the subclass level in CSV format (.csv).

**Supplemental Data 3. Cluster-level markers**. Top 50 robust markers for BICCN cell types at the cluster level in CSV format (.csv).

