## Supplementary figures and images for "How many markers are needed to robustly determine a cell’s type?"

### Supplemental Figures

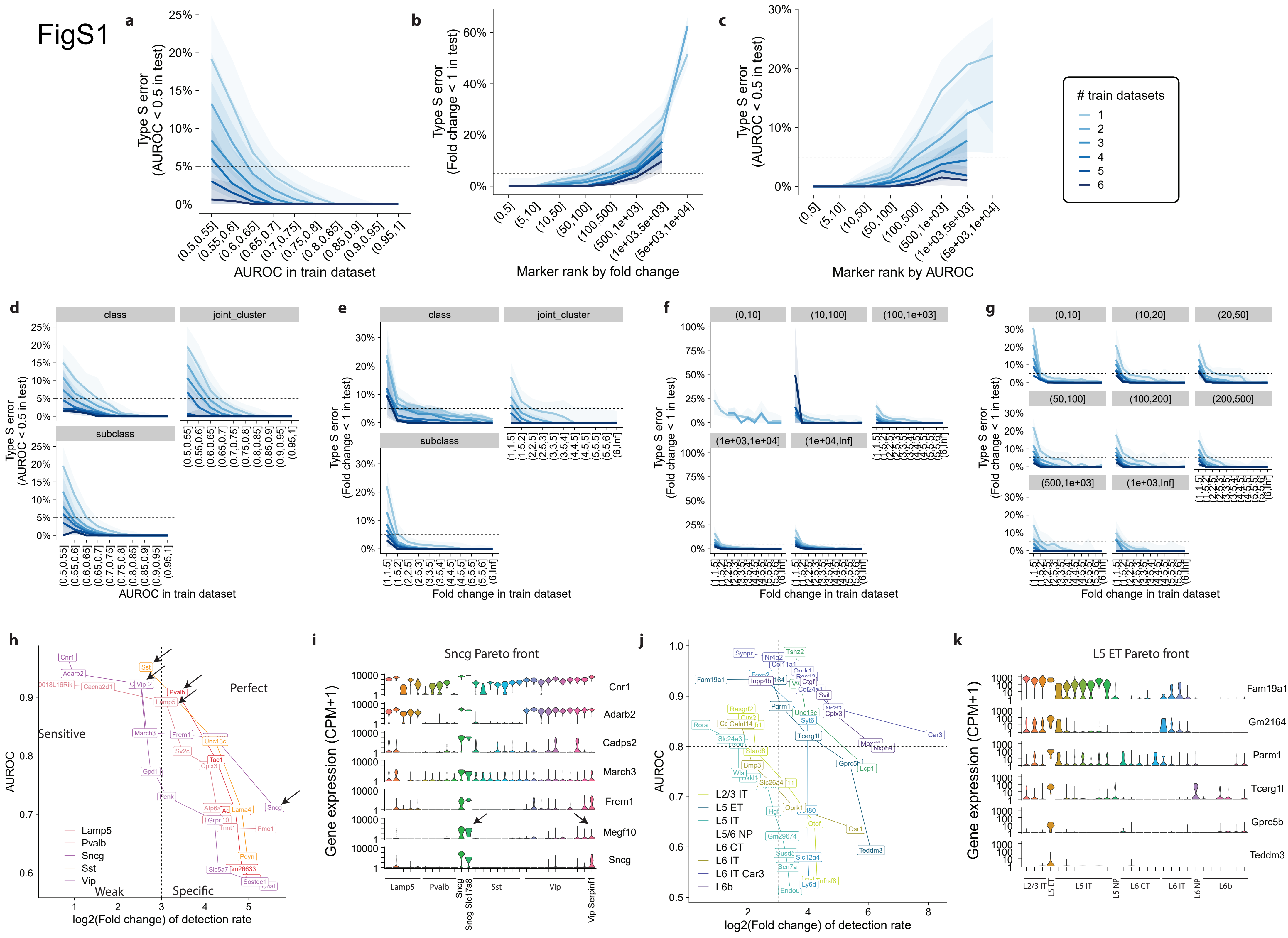

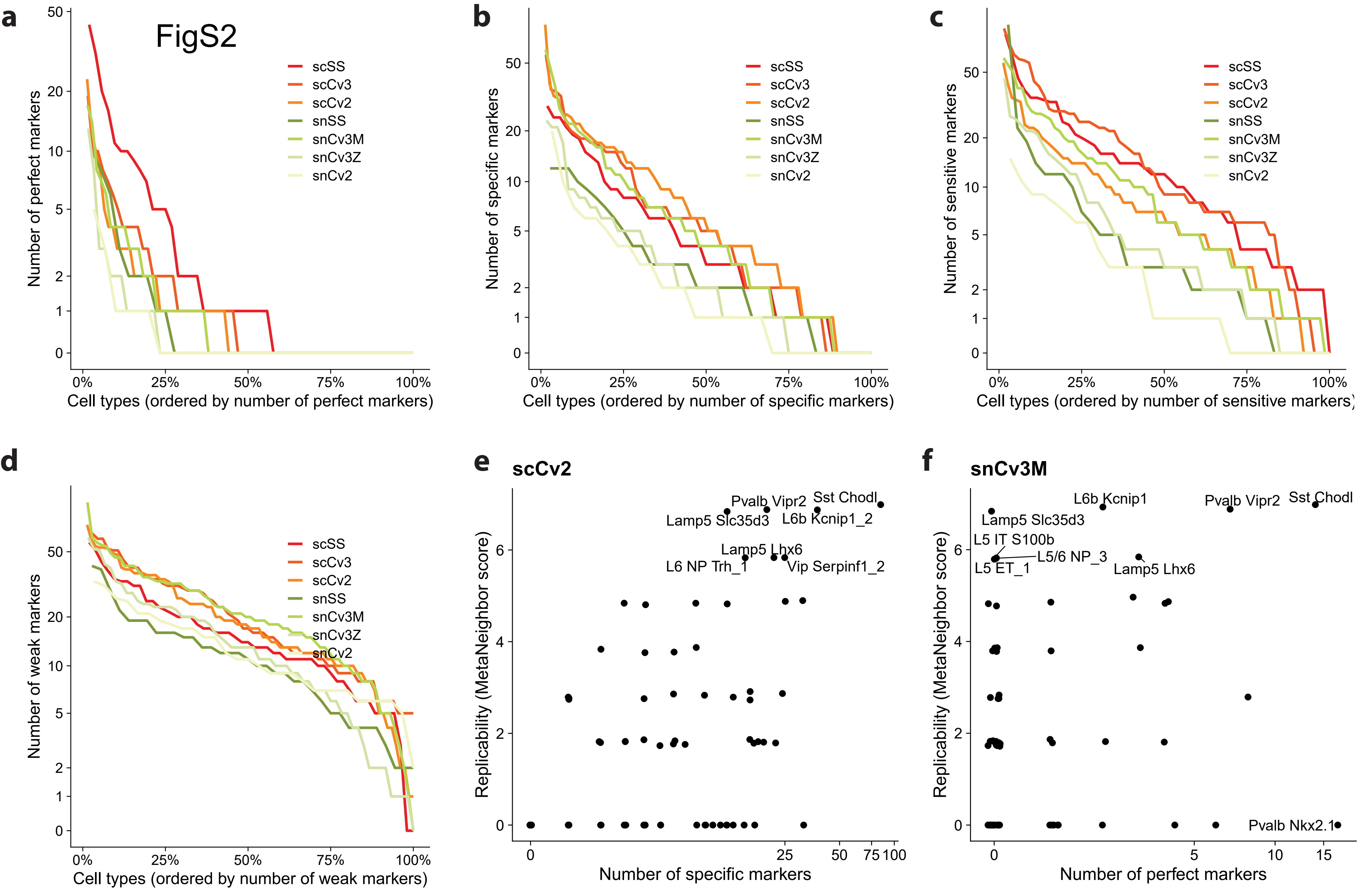

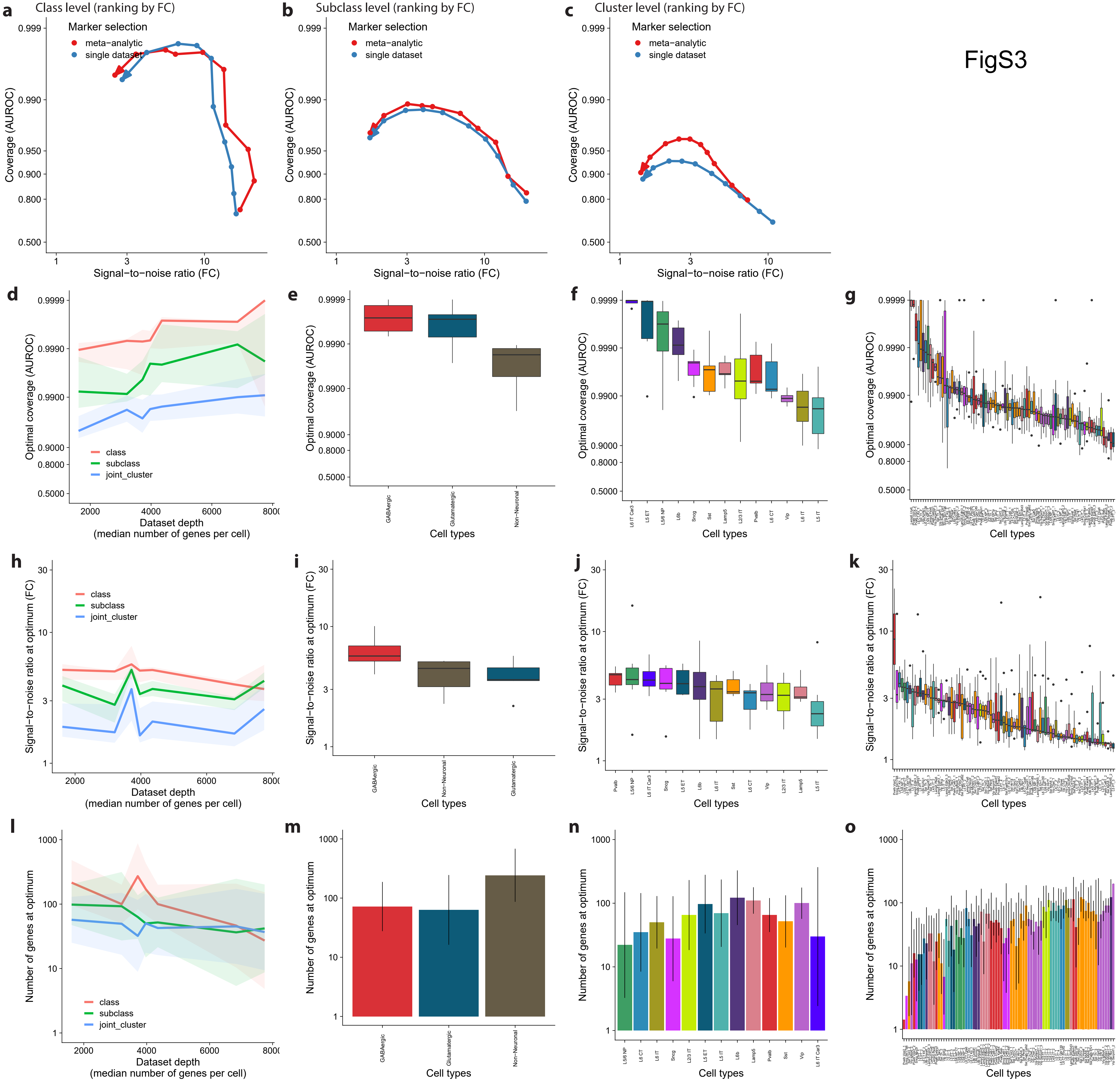

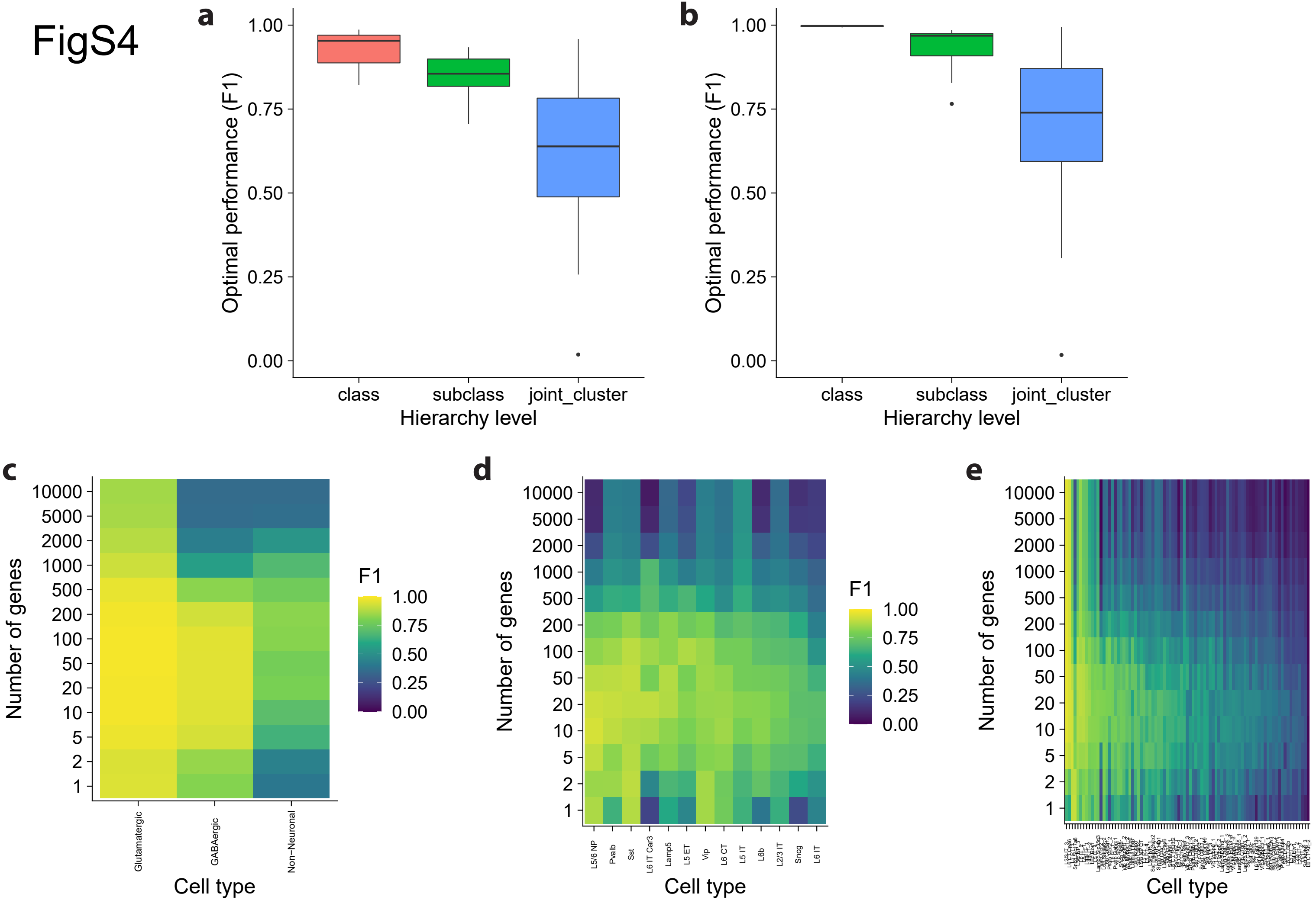

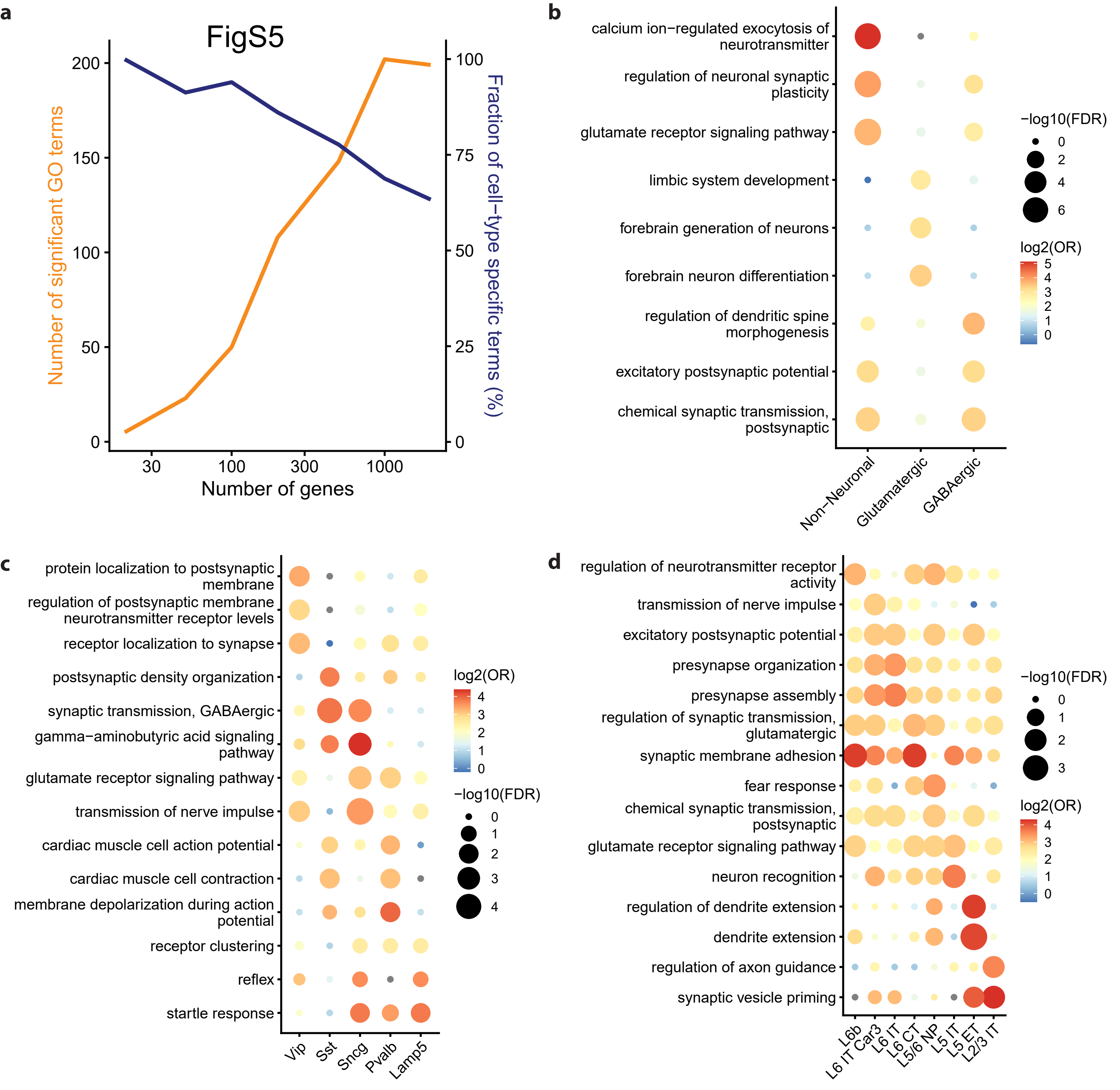

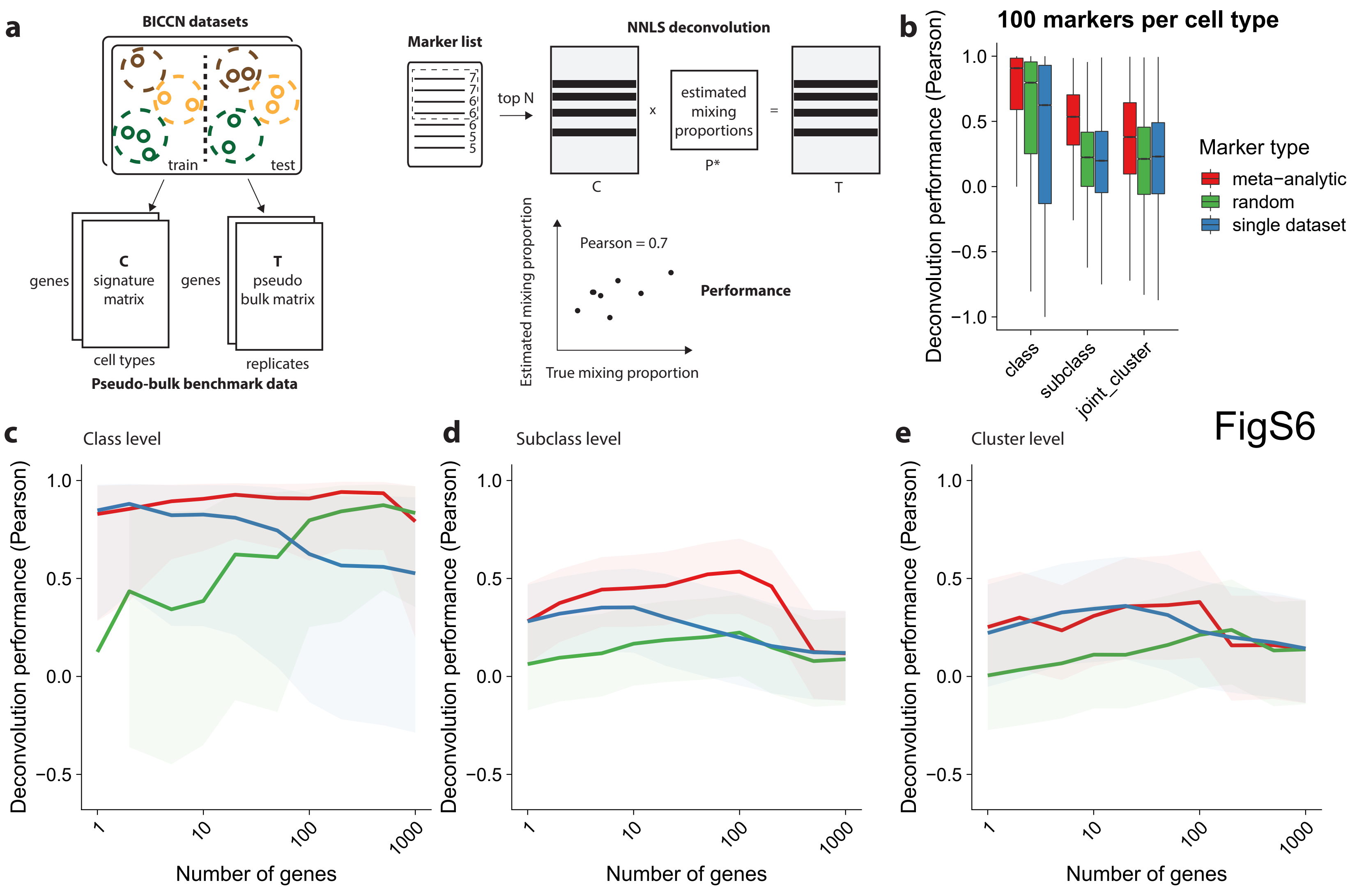

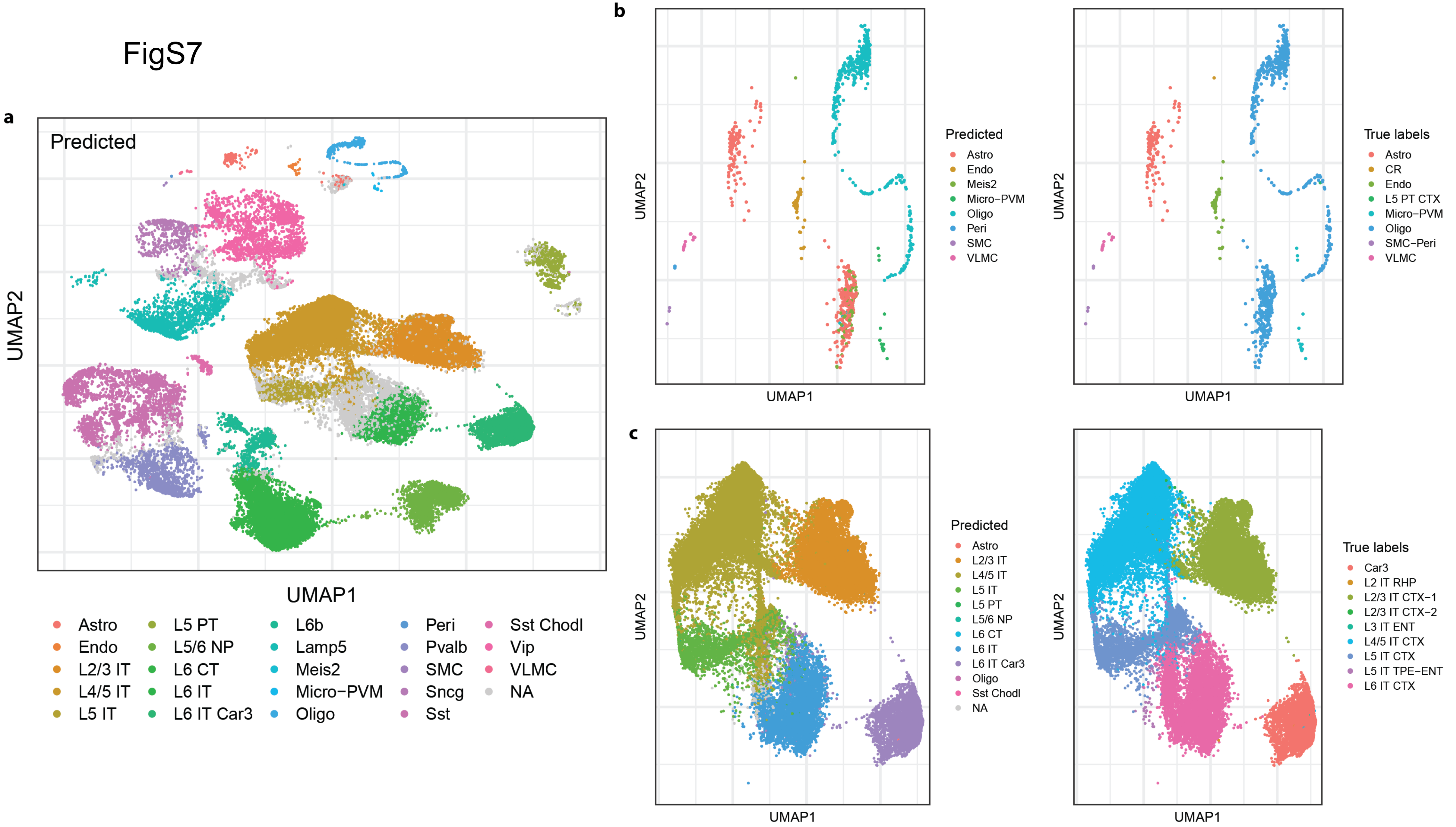

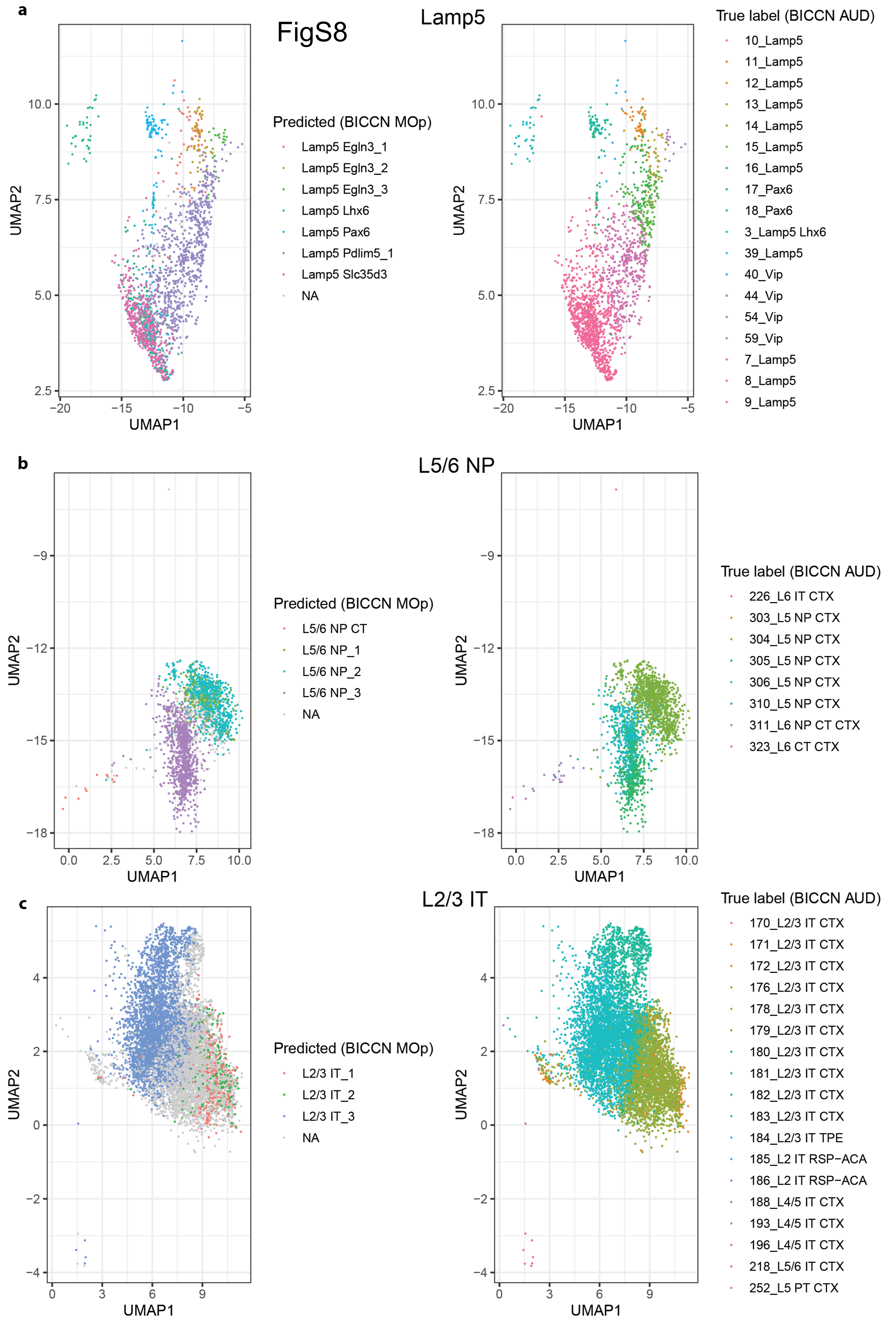

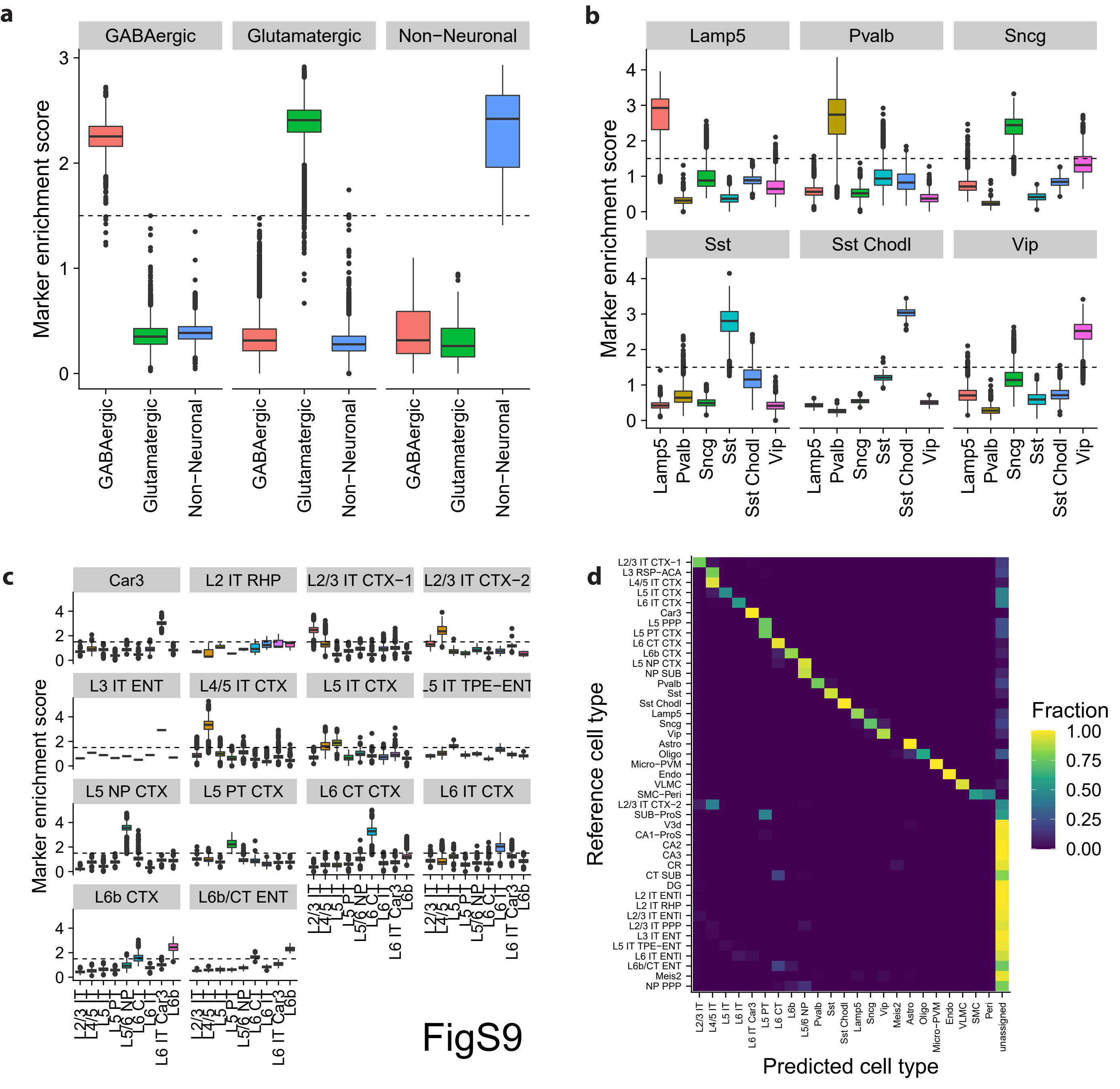
